# Kinome-wide RNAi screen in *Caenorhabditis elegans* reveals new modulators of insulin signaling and longevity

**DOI:** 10.1101/2024.06.10.598287

**Authors:** Manish Chamoli, Anna Foulger, Prachi Singh, Gordon Lithgow, Arnab Mukhopadhyay

## Abstract

The insulin/IGF-I-like signaling (IIS) pathway is a highly conserved signaling cascade that plays a crucial role in regulating longevity across species. Given its significance in aging, identifying novel kinases interacting with the IIS pathway can provide deeper insights into the mechanisms governing longevity. In this study, we performed a targeted RNAi screen of the *Caenorhabditis elegans* kinome, utilizing dauer formation as a phenotypic readout. We identified several known and novel kinase modulators of the IIS pathway. These hits were enriched with both previously documented as well as undocumented lifespan regulators. Thermotolerance assays revealed mixed trends, with some kinase inhibitions increasing while others decreasing protection. We observed a positive correlation between thermotolerance and lifespan extension, as well as between dauer formation and lifespan extension, with thermotolerance proving to be a better predictor of longevity. Our findings offer a valuable resource for researchers exploring the IIS pathway and highlight novel, unannotated kinases as potential new therapeutic targets for aging interventions.

## Introduction

The IIS pathway is a highly conserved signaling cascade that regulates longevity across species (reviewed in^1,2^). Polymorphic variants in FOXO3A, a key downstream effector of the IIS pathway, have been linked to human longevity, underscoring the pathway’s significance in aging processes^3^. Reduced signaling through the IIS pathway can extend the lifespan of *C. elegans* by nearly tenfold^4^. The evolutionary conserved IIS pathway in *C. elegans* involves a kinase-regulated signaling cascade^5,6^, starting with insulin-like ligands binding to the insulin/IGF-1 receptor tyrosine kinase DAF-2. This activation triggers phosphatidylinositol-3-OH kinase (PI3K) AGE-1, subsequently activating PDK-1 (3-phosphoinositide-dependent kinase-1). PDK-1 then phosphorylates and activates the serine/threonine kinases AKT-1 and AKT-2, which inhibit the transcription factor DAF-16, the *C. elegans* homolog of human FOXO, by preventing its nuclear translocation^7,8^. When IIS signaling is reduced, AKT activity decreases, allowing DAF-16 to enter the nucleus and activate genes related to stress resistance, metabolism, and longevity^5,9,10^.

Given the IIS pathway’s extensive influence on various biological functions, identifying novel modulators is essential for gaining a deeper understanding of the mechanisms regulating these processes. Kinases, which constitute a majority of the signaling components within the IIS pathway cascade, are of particular interest. Consequently, we focused on identifying additional kinases that potentially interact with the IIS pathway. To this end, we performed a targeted RNAi screen of the *C. elegans* kinome to identify kinases that affect phenotypes regulated by the IIS pathway. One such phenotype is dauer formation, a developmental stage induced by environmental conditions such as food availability or temperature changes^11,12^. Normally, *C. elegans* larvae pass through four larval (L1, L2, L3 and L4) stages before developing into reproductive young adults. However, under unfavorable conditions, L1 larvae enter a stress-resistant dauer stage^11,12^. The dauer stage represents a form of developmental quiescence, allowing larvae to survive extended periods of unfavorable conditions^13^. This stage is intricately controlled by IIS pathway components. Because of their unique visible appearance, dauers serve as a valuable surrogate phenotype for identifying modulators of the IIS pathway.

In this study, using the dauer phenotype as a readout, we screened the *C. elegans* kinome to identify new kinase modulators of the IIS pathway. We found that the screen hits we identified as modulators are enriched for lifespan regulators and include kinases whose knockdown extends lifespan. By performing a thermotolerance assay on these hits, we show that both dauer formation and thermotolerance positively correlate with lifespan extension, with thermotolerance being a better predictor. Our study provides a valuable resource of new kinases for researchers interested in studying the IIS pathway and novel, unannotated kinases involved in lifespan extension.

## Results

### Kinome-wide RNAi screen for IIS pathway modulators using dauer as the readout

Lack of food causes *C. elegans* larvae to enter a developmental quiescence stage known as dauer^14^. Loss-in-function mutations in *daf-2*, an insulin receptor can also cause larvae to enter dauer stage even in the presence of food^15,16^ (**Fig. 1A**). The partial loss-of-function mutant, *daf-2(e1370)*, can develop into a young adult worm but is highly sensitive to elevated temperatures, entering 100% dauer at 25°C^15,17^. To screen the kinome and identify novel modulators of the IIS pathway, we utilized these hypersensitive mutants.

**Figure 1:**
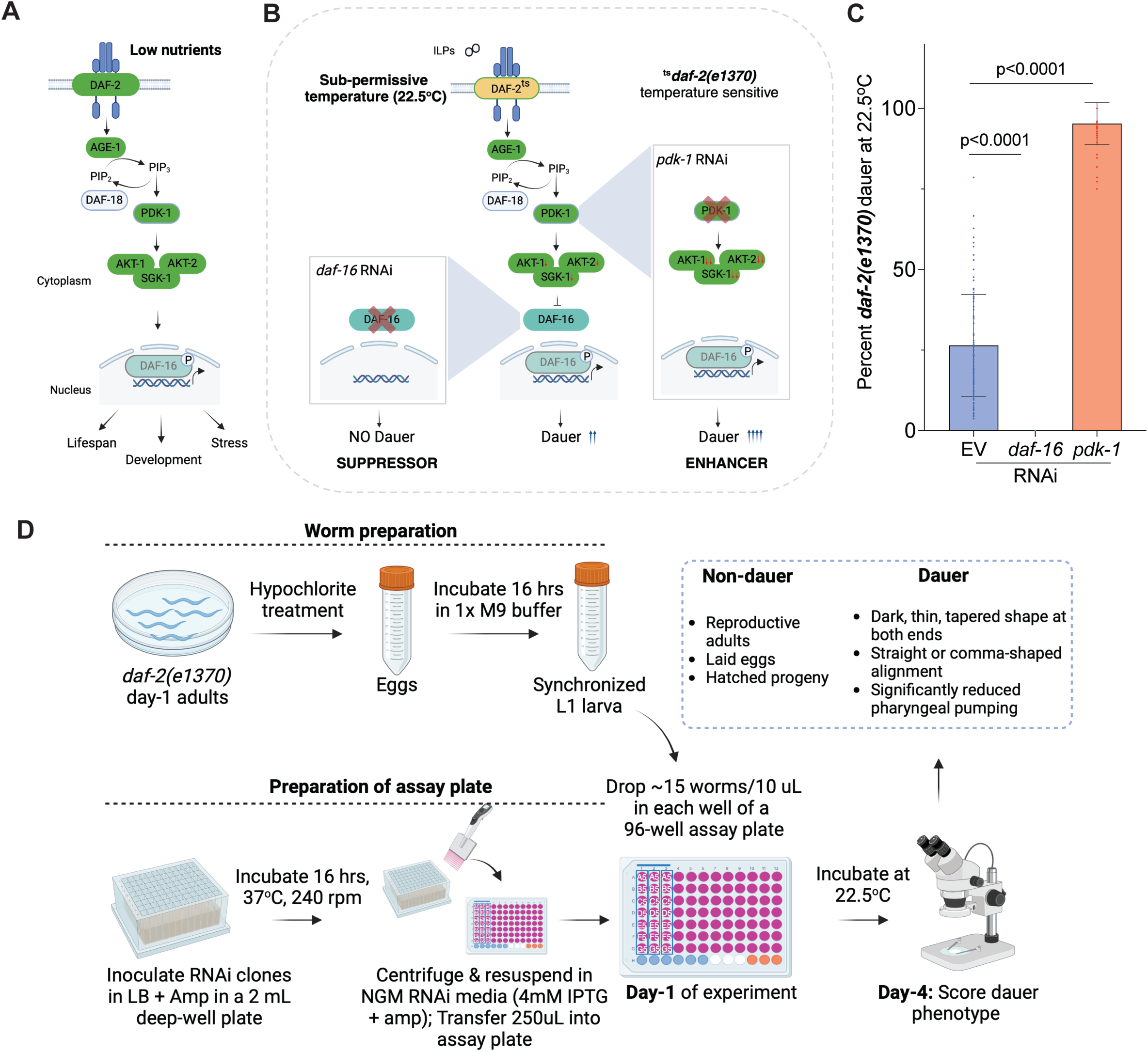
A kinome-wide RNAi screen to identify modulators of the IIS pathway. **(A)** The IIS pathway in *C. elegans* regulates dauer formation. **(B)** Growing temperature sensitive *daf-2(e1370)* mutants at a sub-permissive temperature allows for the identification of both enhancers and suppressors of dauer. **(C)** Percentage of dauer formation (mean ± sd) in empty vector (EV) control (26.46 ± 15.82), *daf-16* (0), and *pdk-1* (95.30 ± 6.50) RNAi-treated *daf-2(e1370)* mutants grown at 22.5°C. p-values were determined using an unpaired Student’s t-test. **(D)** Schematic representation of the kinome-wide RNAi screening workflow.

Our aim was to identify both suppressors and enhancers of the IIS pathway. Thus, our initial experiments were geared towards identifying a sub-permissive temperature that would result in ∼50% dauer (**Fig. 1B**). Under our RNAi screening workflow, we observed that when grown at 22.5°C, *daf-2(e1370)* consistently formed close to 50% dauer. Hence, we decided to conduct our screen at this temperature. We reasoned that for kinases that interact with the IIS pathway, RNAi knockdown would modulate *daf-2(e1370)* dauer formation (**Fig. 1B**). For e.g., RNAi knockdown which will enhance dauer formation are likely to further lower signaling through the IIS pathway, while the one suppressing dauer formation are likely to mediate its downstream effector functions. As a controls in our assays, we used RNAi against *pdk-1* and *daf-16*. The PDK-1 (3-Phosphoinositide-dependent protein kinase-1) is a key kinase in the IIS pathway that activates AKT kinases, essential for inhibiting dauer formation in the presence of food^8^. When PDK-1 activity is reduced by RNAi, IIS signaling is lowered, promoting *daf-2(e1370)* dauer formation. DAF-16, on the other hand, is a FOXO transcription factor that acts downstream of the IIS pathway^18^. DAF-16 is normally inhibited by active IIS signaling i.e., the presence of food, preventing it from entering the nucleus. When IIS signaling is reduced, DAF-16 activates genes related to longevity and dauer formation. Knocking down *daf-16* via RNAi prevents these responses, resulting in reduced dauer formation. These RNAi controls served as positive and negative control, respectively, for the dauer phenotype in our screen (**Fig. 1C**). Our pilot assays with these controls yielded a Z’-factor of 0.79, confirming the scalability of this dauer-based phenotypic assay for kinome-wide RNAi screening.

The screening workflow (**Fig. 1D**; **see methods for detail**) that we used to conduct kinome-wide RNAi screening consisted of 96-wells liquid culture plates, allowing us to screen 31 kinases in triplicate on a single assay plate. Briefly, synchronized cultures of ∼15-20 L1 larvae were dispensed into each well of a 96-well plate containing 60 µL of RNAi bacterial feed. The L1 larvae were allowed to grow for three days at 22.5°C. On the fourth day, when the worms grown on control RNAi or *pdk-1* RNAi entered dauer and *daf-16* RNAi worms became adults, we scored the number of dauers in each well under a light microscope. Dauers were identified by their dark, thin, tapered shape at both ends, aligned straight or in a comma shape with significantly reduced pharyngeal pumping. The percentage of dauers was calculated based on the total number of worms present.

### Kinases identified as modulators of the IIS pathway play roles in diverse biological processes

From the Ahringer RNAi library^19^, which contains over 16,000 RNAi clones, we selected and arranged a mini-96-well kinome RNAi library. Out of the 411 kinase-targeting RNAi clones available in the Ahringer collection, 357 clones produced viable cultures; the rest showed no signs of growth. We used these 357 viable clones to create a curated mini-96-well kinome library. Each plate was formatted to include an empty vector, *pdk-1*, and *daf-16* RNAi, ensuring that each plate contained its respective controls (**Fig. 2A**). Out of the 357 kinases tested, 128 RNAi clones did not exhibit any scorable phenotype. Many of these clones either failed to grow, produced sick worms, or resulted in embryonic lethality. Consequently, we were able to assess the dauer phenotype for only 229 RNAi clones (**Fig. 2B and Supplementary Table 1**). Among these, 75 RNAi clones demonstrated significant changes in dauer formation across biological repeats: RNAi against 69 enhanced, and 6 suppressed *daf-2(e1370)* dauer formation (**Table 1 and Supplementary Table 1**).

**Figure 2:**
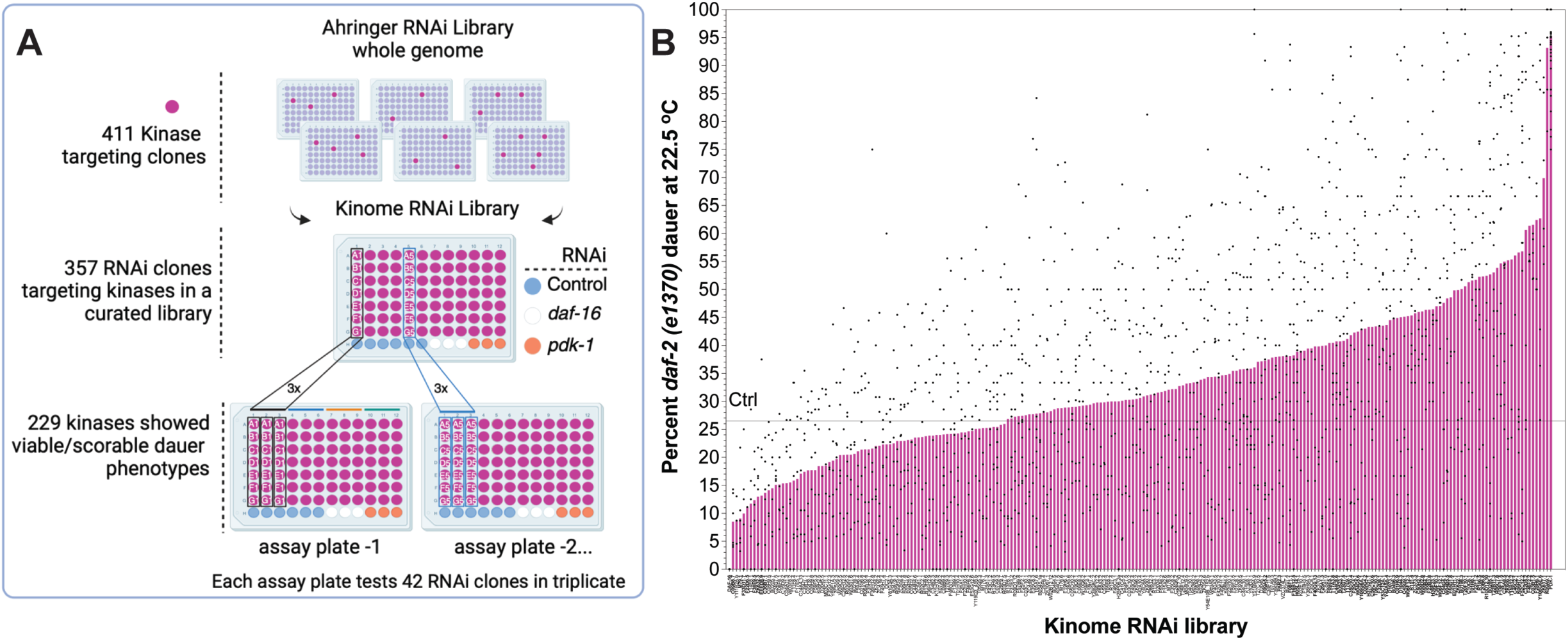
Results of the kinome-wide RNAi screen. **(A)** From the whole genome Ahringer RNAi library, we selected 357 viable kinase-targeting RNAi clones to create a mini-96-well kinome library plates. Each plate included controls: an empty vector control, *daf-16* and *pdk-1* RNAi. Out of 357 clones, 229 RNAi clones showed scorable dauer phenotype. **(B)** Graphical representation of all 229 kinase RNAi clones showing enhancers and suppressors of *daf-2(e1370)* dauer. Bars represent the mean, and each dot in the graph represents an individual well with 15-20 worms across two independent trials. See Table 1 and Supplementary Table 1 for more details.

**Table 1:**
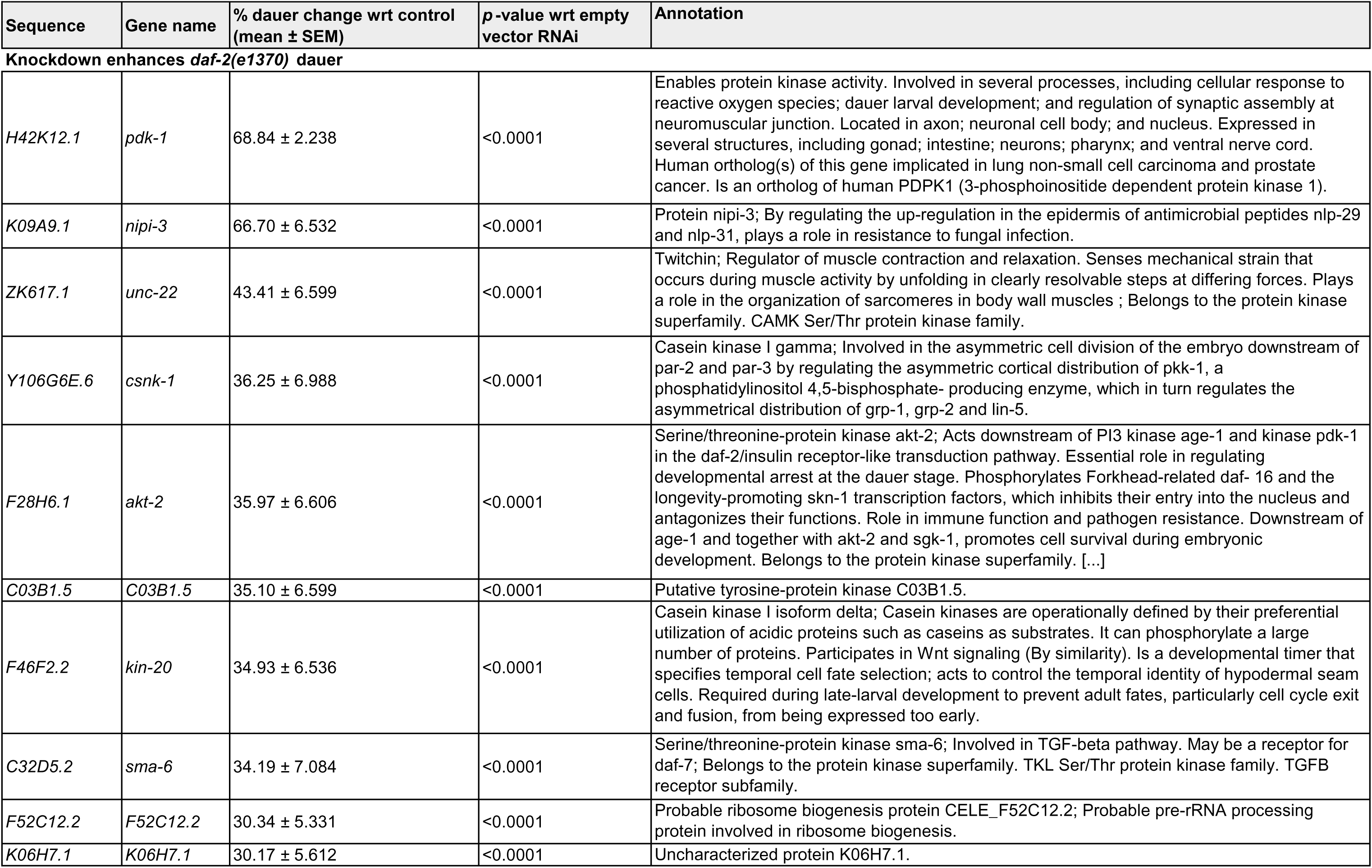
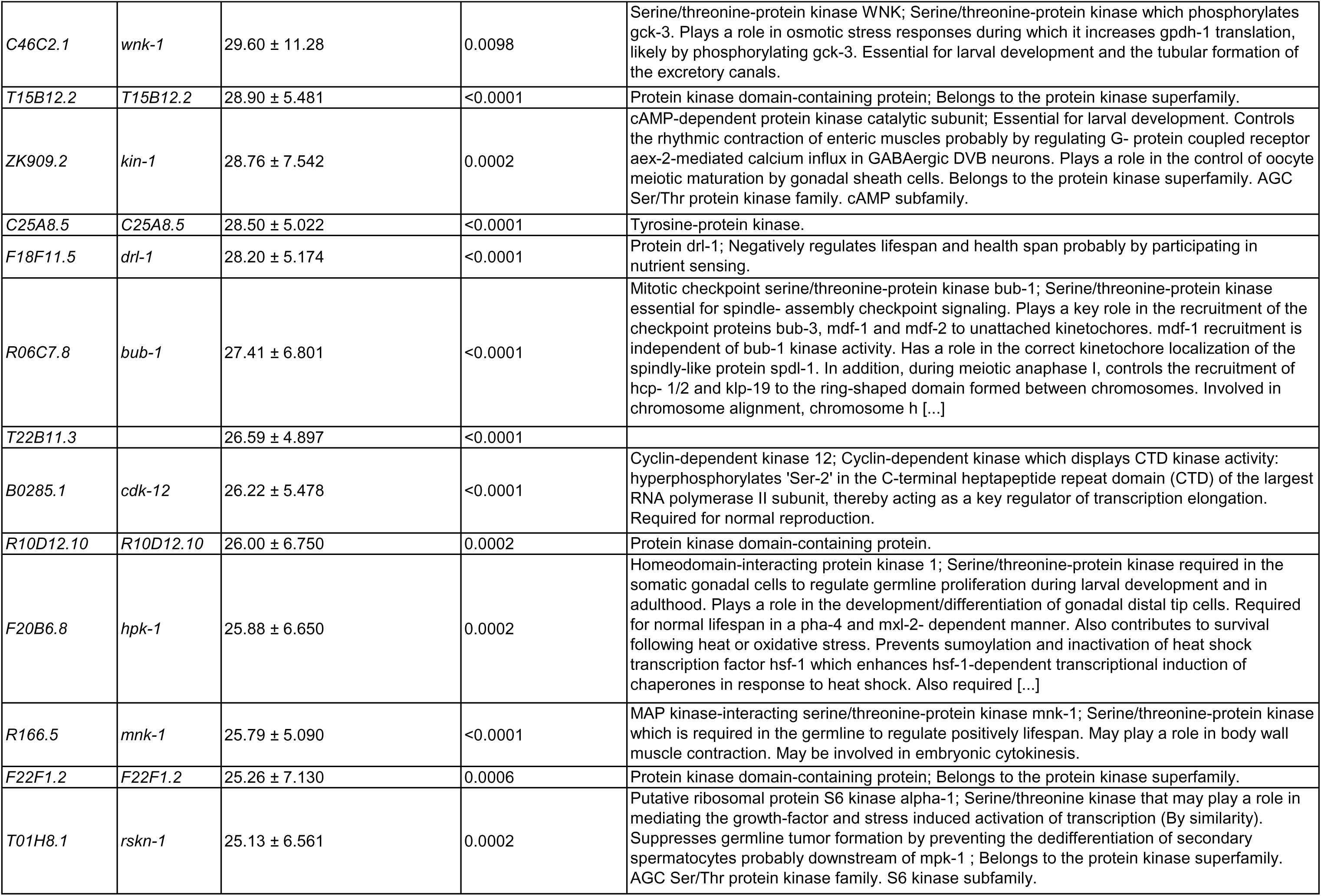
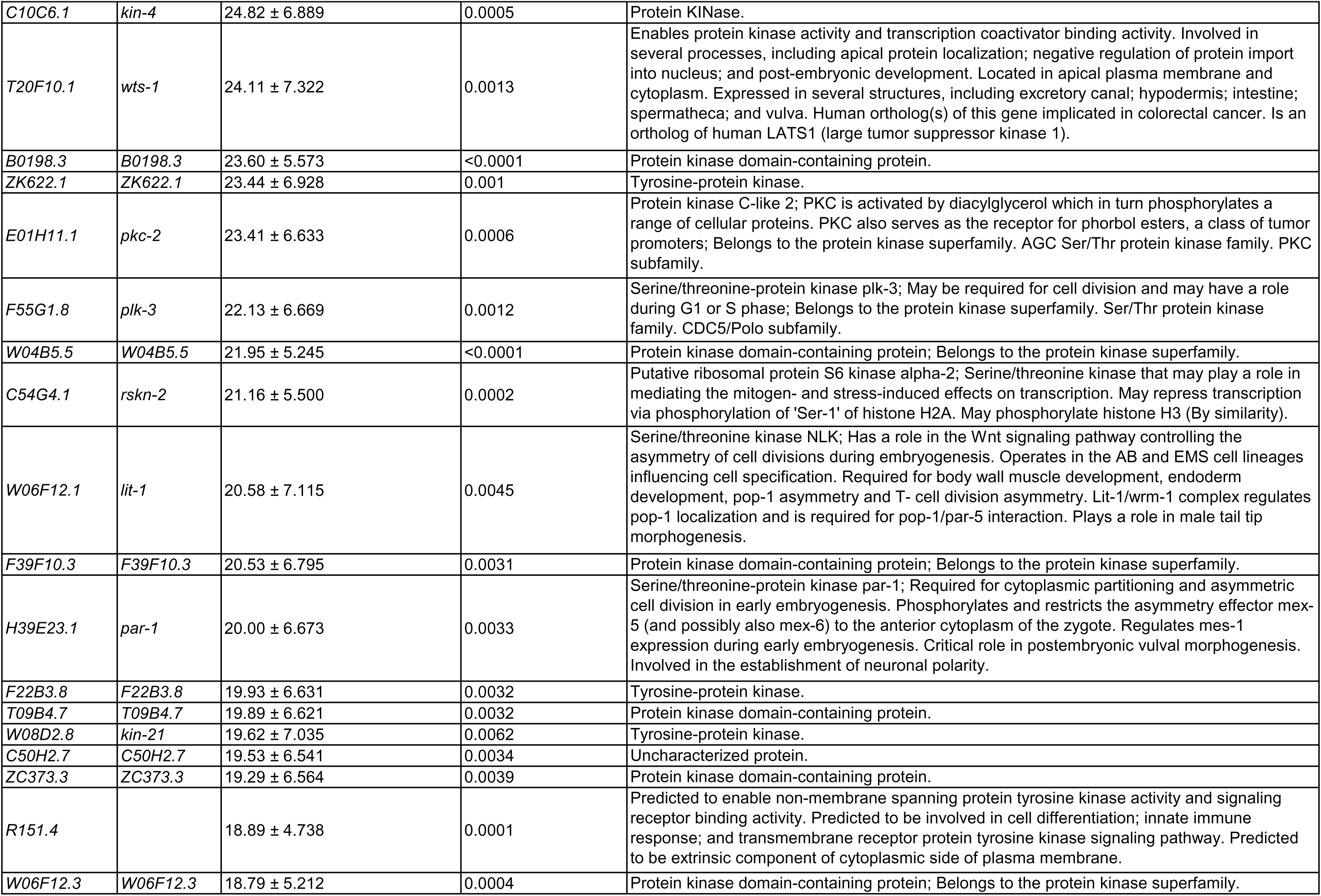
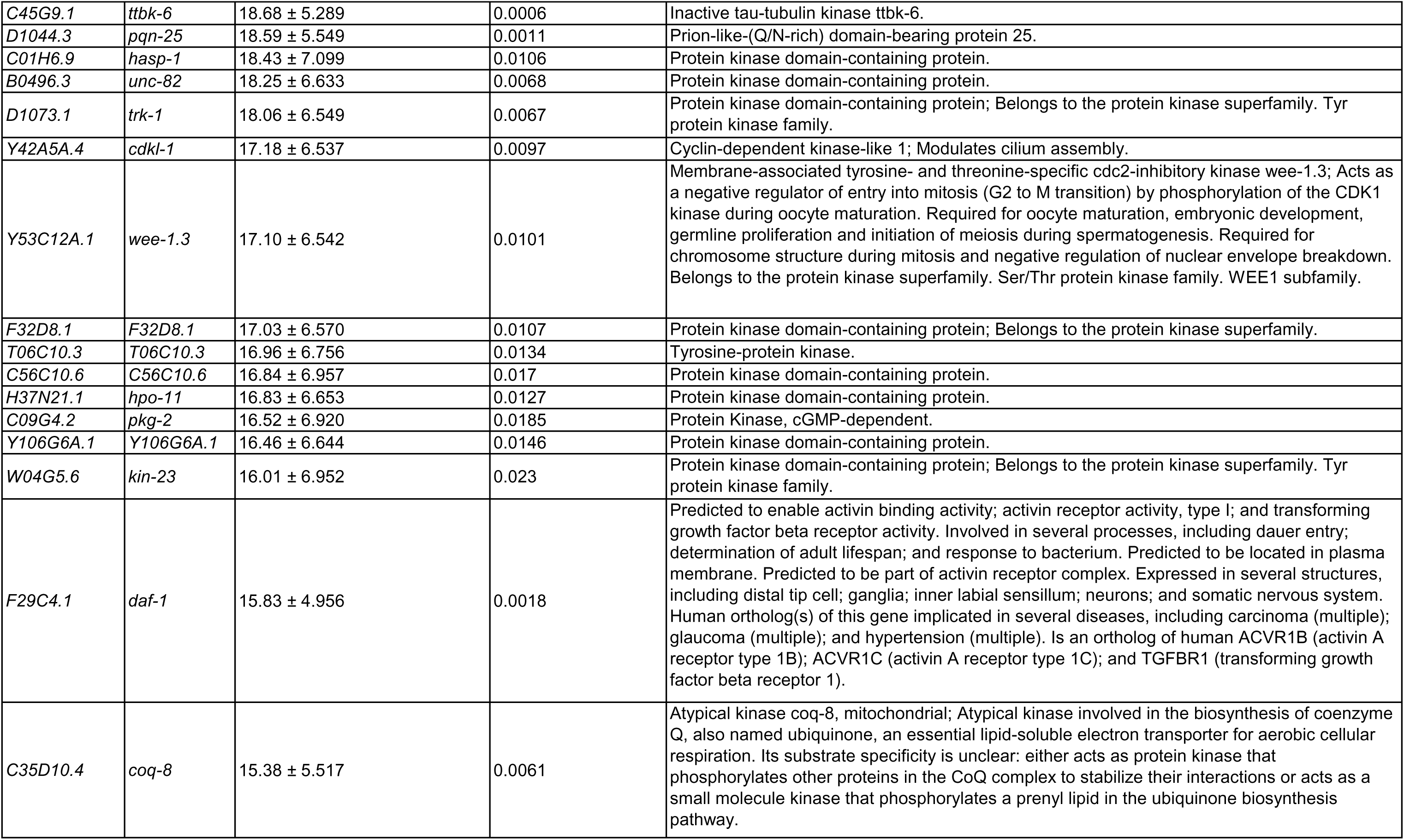
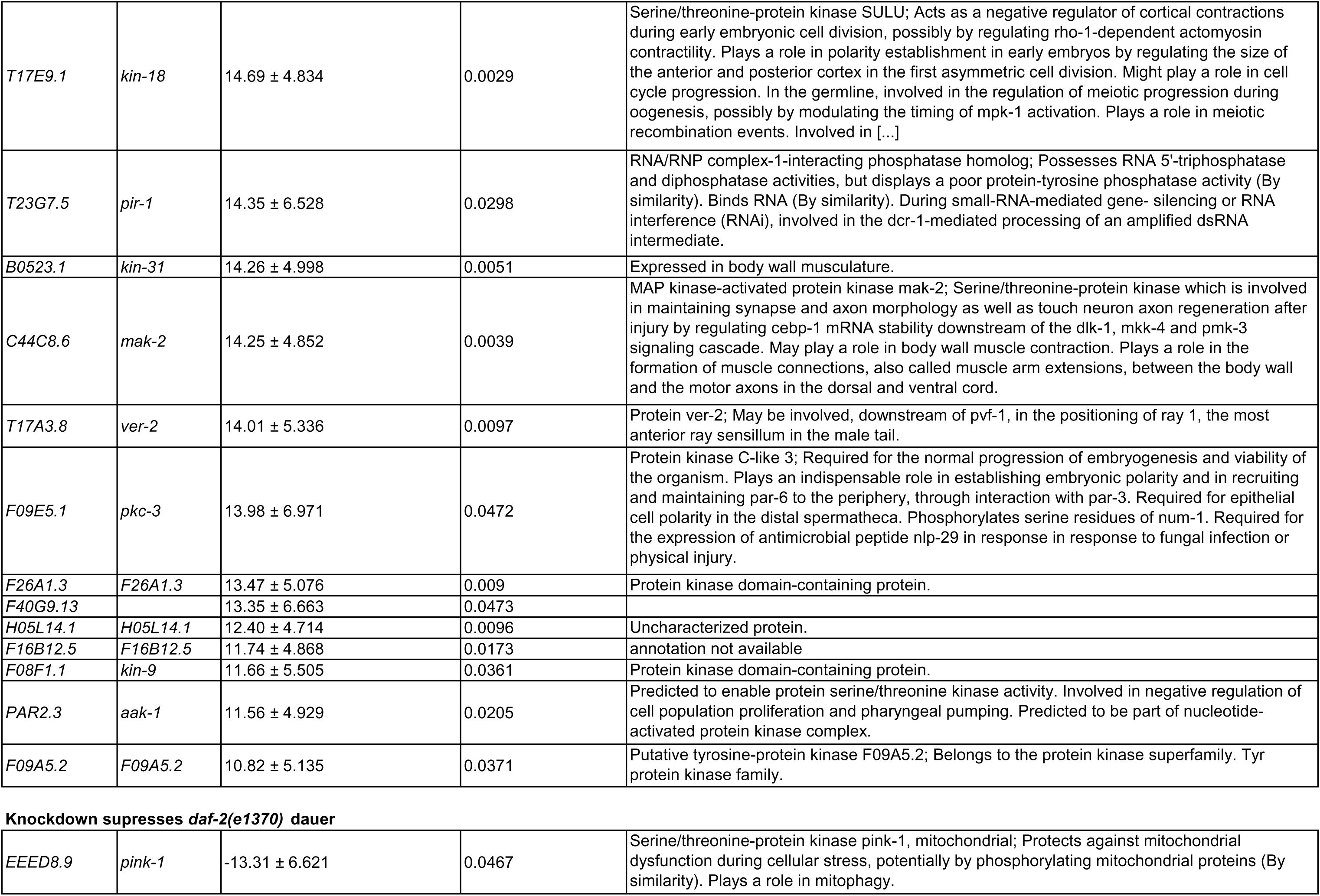
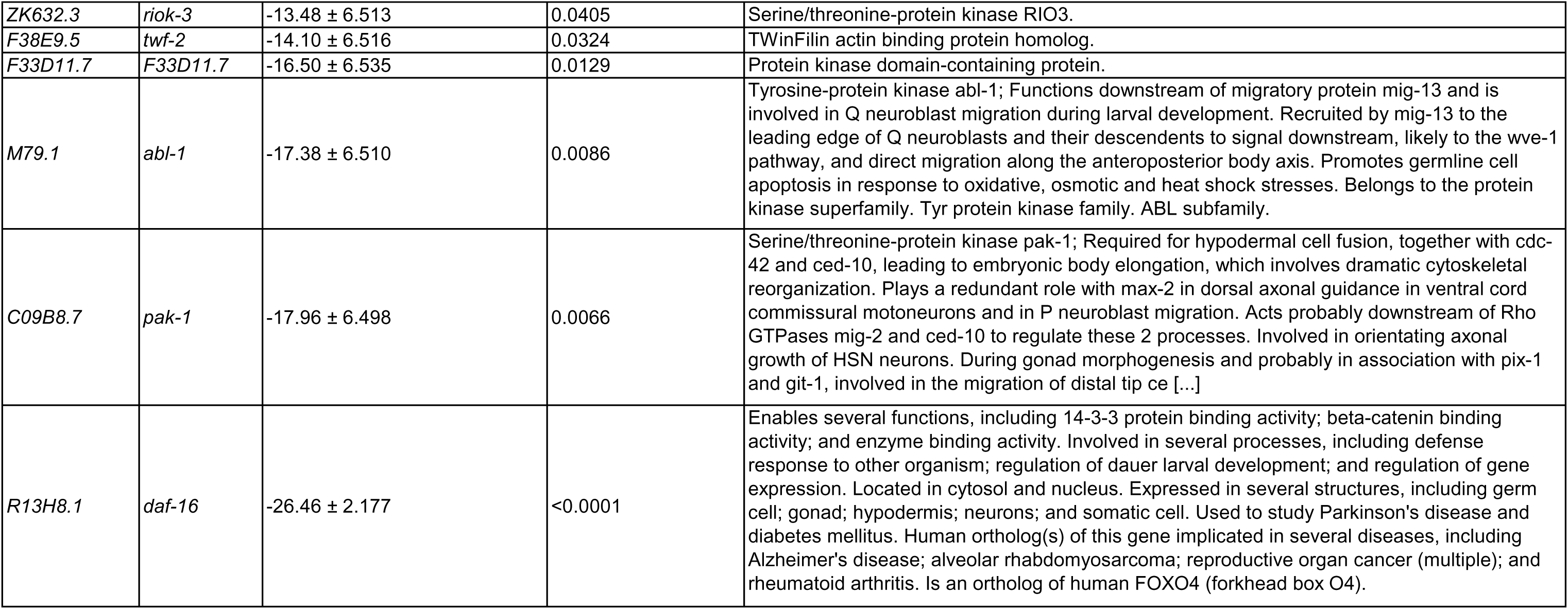
Enhancer and suppressor of *daf-2(e1370)* dauer. List of hits and their annotations from the kinome RNAi screen that, when knocked down, enhance or suppress *daf-2(e1370)* dauer formation at 22.5°C. The percentage change in dauer formation is shown relative to the empty vector control RNAi, with significance determined using an unpaired Student’s t-test. See Supplementary Table 1 for the percentage of dauer formation from all the kinases tested.

The screen yielded a broad list of kinases with diverse regulatory roles across various biological processes, emphasizing their roles in cellular signaling, development, and stress responses. The identification of several hits known to be involved in the IIS pathway and TGF-beta pathway highlights the success of the screen, which was designed to identify modulators of the IIS pathway **(Table 1 and Supplementary Table 1)**. One notable example is serine/threonine-protein kinase AKT-2, which functions downstream of PI3 kinase AGE-1 and kinase PDK-1 within the DAF-2/IIS pathway^8,20^. This kinase plays a central role in regulating developmental arrest at the dauer stage. In the context of stress response, the serine/threonine-protein kinase WNK-1 is significant for its role in osmotic stress responses, where it increases GPDH-1 translation by phosphorylating GCK-3^21,22^. Additionally, serine/threonine-protein kinase SMA-6 is a critical component of the TGF-beta signaling pathway, potentially acting as a receptor for DAF-7 and contributing to various developmental processes^23,24^. The TGF-beta signaling pathway is also a known regulator of reproductive longevity and innate immune response^24^.

The screen also identified kinases known to regulate lifespan, including DAF-1 (TGF-beta receptor type I), which plays a crucial role in the TGF-beta signaling pathway affecting dauer formation^25^. HPK-1 (homeodomain-interacting protein kinase 1) is involved in stress response pathways, influencing longevity^26,27^. MNK-1 (MAP kinase-interacting serine/threonine-protein kinase 1) contributes to lifespan regulation through its roles in muscle function and MAP kinase interactions^28^. WTS-1 (Warts, also known as large tumor suppressor kinase 1), part of the Hippo signaling pathway, impacts cell proliferation and lifespan^26^. PAK-1 (p21-activated kinase 1) modulates cytoskeletal dynamics and cell signaling, affecting aging^29,30^.

Cell-cycle regulation is prominently represented among the identified kinases. Notably, mitotic checkpoint serine/threonine-protein kinase BUB-1 is essential for spindle-assembly checkpoint signaling and chromosome alignment during cell division, ensuring proper genomic stability^31,32^. Additionally, Cyclin-dependent kinase 12 is crucial for transcription regulation during the cell cycle, displaying CTD kinase activity is necessary for transcription elongation^33^. Furthermore, membrane-associated tyrosine- and threonine-specific CDC2-inhibitory kinase WEE-1.3 acts as a negative regulator of the G2 to M transition by phosphorylating CDK1, thereby controlling entry into mitosis and playing critical roles in oocyte maturation, embryonic development, and chromosome structure during mitosis^34–36^.

We also identified kinases with roles in immune defense mechanisms, such as NIPI-3 and Protein kinase C-like 3 (PKC-3), which play crucial roles in regulating antimicrobial peptides and enhancing resistance to fungal infections^37,38^. NIPI-3 regulates the upregulation of NLP-29 and NLP-31 in the epidermis^39^, while PKC-3 is required for the expression of NLP-29 in response to fungal infection or physical injury^38^.

Overall, our kinome-wide RNAi screen revealed a diverse array of kinases involved in critical biological processes, including the IIS and TGF-beta pathways, cell-cycle regulation, and immune defense.

### Discovery of new kinase regulators of longevity

Given the well-established role of the IIS pathway in regulating longevity, we were keen to explore how RNAi knockdown of these kinases could influence lifespan. To identify novel longevity regulators, we performed a lifespan analysis on the hits identified from our kinome RNAi screening, utilizing a modified version of the 96-well lock-in assay for high-throughput lifespan analysis^40^ (see Methods for detail). From this initial screening, we identified 31 RNAi clones as positive hits (**Supplementary Table 2 and Supplementary Table 4**). Subsequently, we tested these clones on NGM bacterial RNAi plates. Although all the hits showed a trend toward increased lifespan when knocked down post-developmentally, only 14 RNAi clones were found to significantly extend lifespan (**Fig. 3A and 3B**, **Table 2 and Supplementary Table 2**).

**Figure 3:**
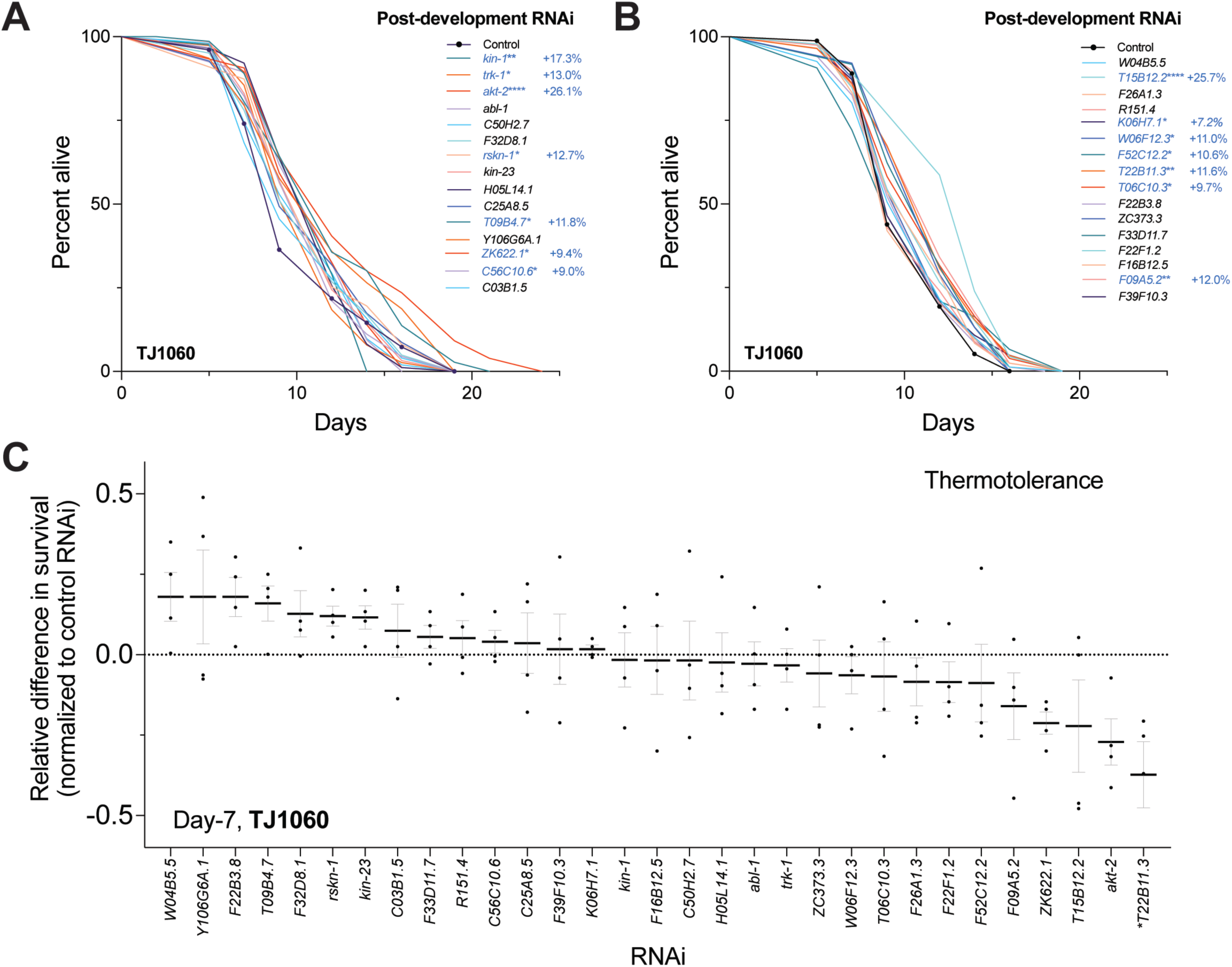
Lifespan and thermotolerance assays of hits from the kinome-wide RNAi screen. **(A)** Lifespan graph of 31 hits validated on NGM RNAi plates. RNAi clones that significantly increase lifespan are colored in blue, with p-values indicated as follows: *p<0.05, **p<0.01, and ****p<0.0001. p-value determined using Mantel-Cox log rank test. See Table 2 and Supplementary Table 2 for more details on percent changes. **(B)** Relative difference in survival normalized to empty vector control RNAi of day-7 TJ1060 worms post-exposure to 35°C for 12 hours. p-values were determined using an unpaired Student’s t-test, *p<0.05. See Supplementary Table 3 for more details on percent changes across trials

**Table 2:** List of kinases and their human orthologs, with descriptions, that significantly extend lifespan when knocked down. Strain: TJ1060. Temperature: 25°C. Post-developmental RNAi treatment. See Figure 3A, 3B, and Supplementary Table 2 for details.

**Knockdown significantly extends lifespan**
| **Sequence** | **Gene name** | **Human Ortholog** | **Description (ortholog)** |
| --- | --- | --- | --- |
| *ZK909.2* | *kin-1* | *PRKACB* | protein kinase cAMP-activated catalytic subunit beta |
| *D1073.1* | *trk-1* | *NTRK3* | neurotrophic receptor tyrosine kinase 3 |
| *F28H6.1* | *akt-2* | *AKT/PKB* | Protein kinase B/AKT |
| *T01H8.1* | *rskn-1* | *RPS6KA3* | ribosomal protein S6 kinase A3 |
| *ZK622.1* | *ZK622.1* | *FER/FES* | FER tyrosine kinase |
| *C56C10.6* | *C56C10.6* | *TTBK2* | tau tubulin kinase 2 |
| *T15B12.2* | *T15B12.2* | *CSNK1A1* | casein kinase 1 alpha 1 |
| *W06F12.3* | *W06F12.3* | *TTBK2* | tau tubulin kinase 2 |
| *F52C12.2* | *F52C12.2* | *TSR3* | TSR3 ribosome maturation factor |
| *T22B11.3* |  | *LTK* | leukocyte receptor tyrosine kinase |
| *T06C10.3* | *T06C10.3* | *FER* | FER tyrosine kinase |
| *F09A5.2* | *F09A5.2* | *TIE1* | tyrosine kinase with immunoglobulin like and EGF like domains 1 |
| *K06H7.1* | *K06H7.1* |  |  |
| *T09B4.7* | *T09B4.7* |  |  |

Strain: TJ1060; Temperature: 25C; post-development RNAi
See Supplementary. Table 2 for lifespan stats

Out of the 14 RNAi treatments that significantly extended lifespan, only four have been annotated: *kin-1*, *trk-1*, *rskn-1* and *akt-2*. The other eight, namely *ZK622.1*, *C56C10.6*, *T15B12.2*, *W06F12.3*, *F52C12.2*, *T22B11.3*, *T06C10.3*, *F09A5.2*, *K06H7.1* and *T09B4.7*, remain unannotated with no available information on their functions. *kin-1*/PKA has been shown to play a role in the innate immune response and protection against cold-induced stress^41,42^. *trk-1*/TRK is a homolog of the neurotrophin receptor tyrosine kinase in *C. elegans* and a substrate for the LAR family protein tyrosine phosphatase *ptp-3*^43^. *akt-2*/AKT is involved in the IIS pathway, directly contributing to the sequestration of DAF-16 in the cytoplasm^20^. The human ortholog of unannotated kinases we identified have been reported to play role in important cellular processes and disease mechanisms. For instance, C56C10.6/ TTBK2, Tau Tubulin Kinase 2 is intriguing due to its involvement in phosphorylating tau proteins, which are implicated in neurodegenerative diseases like Alzheimer’s^44,45^. Similarly, T15B12.2/ CSNK1A1 Casein Kinase 1 Alpha 1 is known for its role in regulating key signaling pathways that influence circadian rhythms and cellular responses to DNA damage^46^. Another protein of interest is the F52C12.2/TSR3, Ribosome Maturation Factor, essential for ribosome biogenesis and therefore, important in regulating protein synthesis and cellular growth^47^. The T22B11.3/LTK, Leukocyte Receptor Tyrosine Kinase, primarily expressed in hematopoietic tissues, plays a critical role in immune response modulation and development^48^. Further investigating these kinases in the context of their role in aging could provide valuable insights into how these crucial processes influence longevity and age-related diseases.

Overall, our lifespan analysis indicates that several kinases identified by the dauer phenotypic screen may regulate lifespan, showing increased effects when knocked down post-developmentally. These findings merit further investigation to elucidate their roles in aging processes.

### Correlation of thermotolerance and dauer with lifespan extension in IIS pathway modulators

To assess whether the increased lifespan observed following the knockdown of specific kinases is accompanied by enhanced healthspan, we evaluated their sensitivity to an external stressor. Typically, an organism’s ability to withstand noxious temperatures—a measure known as thermal stress resistance—declines with age^49^. In our experiments, we noted a significant decline in the survival rate of the wild-type sterile strain exposed to elevated temperatures (35°C) for 12 hours; survival decreased from 78% on day 6 to 38% on day 9 in aged worms (**Supplementary Fig. 1**). Hence, we decided to determine how the 31 positive primary hits identified in our study influence thermotolerance. We conducted thermotolerance assays on day 7 of adulthood, following exposure to respective RNAi from day 1. Control worms treated with an empty vector exhibited approximately 50% survival after 12 hours at 35°C. Although we observed mixed trends toward both increased and decreased survival after thermal stress, none of these changes reached statistical significance, except *T22B11.3* (**Fig. 3C, Supplementary Table 3)**. This is likely due to underlying variability in the thermotolerance assay^50^. Nonetheless, a correlation analysis examining the relationships between lifespan and thermotolerance, as well as lifespan and dauer, across each RNAi treatment, revealed significant positive correlations. We observed a Pearson correlation coefficient (r) of 0.56 (**Fig. 4A**; R²=0.31, p=0.0009), indicating that thermotolerance is a better predictor of lifespan extension compared to dauer formation, which showed a Pearson r of 0.45 (**Fig. 4B**; R²=0.21, p=0.009). This suggests that, despite experimental variability in thermotolerance, thermal resistance may serve as a more reliable proxy for identifying genes that extend lifespan—a finding that aligns with a recent study identifying thermotolerance assays as better predictor for screening lifespan-extending interventions^50^.

**Figure 4:**
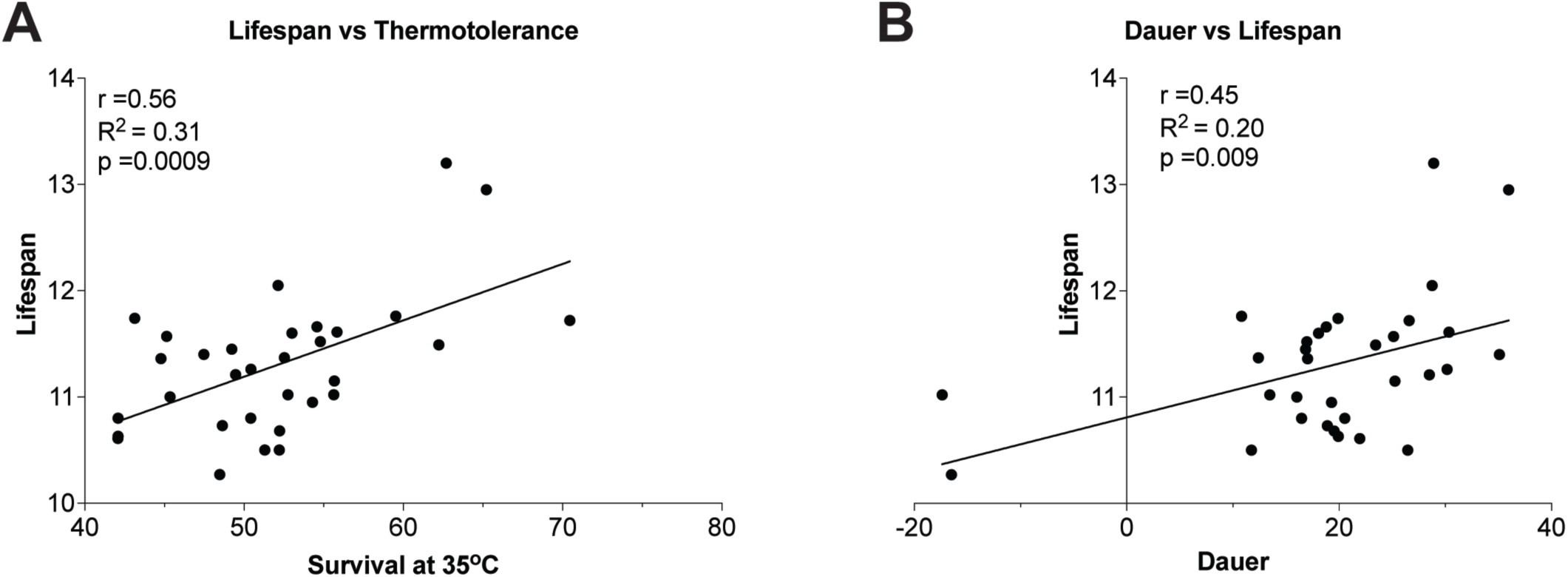
Correlation of thermotolerance and dauer with lifespan extension. Kaplan-Meier means were used for lifespan, and means were used for thermotolerance and dauer. **(A)** Lifespan vs. Thermotolerance; Pearson correlation coefficient (r) = 0.56, R² = 0.31, p = 0.0009. **(B)** Dauer vs. Lifespan; Pearson r = 0.45, R² = 0.21, p = 0.009. P-values were determined using a two-tailed test.

## Discussion

In this study, we screened for novel kinase modulators of the IIS pathway using *C. elegans* dauer formation as a phenotypic readout. Utilizing the hypersensitive *daf-2(e1370)* mutant and a feeding RNAi library, we performed RNAi knockdown of 81% of the *C. elegans* kinome. We were able to quantify dauer phenotypic scoring for approximately 64% of the kinome tested, as the remaining 36% showed no quantifiable phenotype due to larval lethality and arrest. From this, we identified 75 kinases that significantly affected dauer formation when inhibited by RNAi. Notably, 69 RNAi treatments enhanced dauer formation, while 6 suppressed.

The screen revealed a broad list of kinases with diverse regulatory roles, including those in the IIS and TGF-beta pathways, cell-cycle regulation, and immune defense. Our lifespan analysis of these hits revealed that inhibiting the activity of 14 kinases significantly extended lifespan, including *kin-1* and *trk-1*, along with ten unannotated kinases. This indicates that these kinases may play critical roles in modulating lifespan through their interaction with the IIS pathway. Furthermore, a positive correlation between thermotolerance and increased lifespan suggests that thermotolerance could serve as a reliable predictor for longevity interventions. These findings provide a foundation for further exploration of the identified kinases in aging and stress signaling, potentially leading to novel therapeutic targets.

While our lifespan screen identified only a few subsets of kinases whose knockdown significantly increased lifespan in wild type, we observed a general trend toward increased lifespan for many of the kinases tested, with ∼40% of hits showing positive trends. This suggests that hits from the screen are enriched for lifespan regulators. We performed our lifespan assay at 25°C in a sterile strain background to avoid the confounding effects of fluorodeoxyuridine (FUdR)^51,52^, with RNAi knockdown initiated post-developmentally. We chose to conduct post-developmental RNAi knockdown to avoid any developmental effects caused by kinase knockdown. Therefore, it is worth testing these dauer-modulating hits under different conditions where RNAi knockdown is initiated from the larval stage onwards. For instance, over the past years, we have extensively characterized one of the hits from this screen, DRL-1^53,54^, and its homolog FLR-4^55^. DRL-1 kinase knockdown extends mean lifespan by nearly 60% when initiated at the L1 larval stage but fails to extend lifespan when initiated post-developmentally^53^. On the other hand, FLR-4 is unique as its knockdown extends lifespan only under specific bacterial diets^55^.

An important next step will be to test these kinase hits for their specific roles in mediating phenotypes other than longevity associated with the long-lived IIS pathway mutants like *daf-2*. In this direction, our recent study with one such identified kinase hit, B0285.1/CDK-12, shows that it is crucial for maintaining DNA integrity. We used CDK-12 as a genetic tool to analyze the effects of tissue-specific DDR perturbation and DNA damage in the IIS pathway mutants^56^. We found that knocking down *cdk-12* only in the somatic uterine tissue of the IIS pathway mutants led to a DAF-16-dependent pachytene arrest of germ cells. This was achieved through the lowering of the ERK MAPK pathway below a critical threshold.

Together, our study identifies a diverse set of novel kinases that interact with the IIS pathway to regulate its functional outputs.

## Methods

### *Caenorhabditis* elegans strains and maintenance

*C. elegans* strains used in the study are CB1370: *daf-2(e1370) III* and TJ1060: *spe-9(hc88) I; rrf-3(b26) II*. The strains were obtained from the *Caenorhabditis* Genetics Center (CGC), University of Minnesota, MN, USA. Strains were maintained at 15 °C under standard laboratory conditions, as described previously^57^. Worm populations were maintained in 60 mm NGM agar plates (3 g/L NaCl, 17 g/L agar; 2.5 g/L peptone; 1 mM CaCl2, 5 mg/L cholesterol, 1 mM MgSO4, 25 mM KPO4) seeded with OP50 *Escherichia coli*.

### Kinome-wide RNAi Screen

**Day 0**

**Inoculation of RNAi bacterial stock**

1. 500 µl of LB media containing ampicillin (100 µg/ml) was dispensed into a 2 ml 96 deep-well plate using a multichannel pipette (50–1200 µl).

2. The LB was inoculated with frozen RNAi bacterial stock using a multichannel pipette (5–100 µl).

3. The deep-well culture plates were sealed with a breathable air-sealer to allow air exchange and prevent contamination.

4. For bacterial growth, these culture plates were placed in a shaking incubator at 240 rpm, 37°C for 16 hours.

**Sodium hypochlorite treatment for L1 larvae synchronization**

5. The *daf-2(e1370)* mutants were propagated at 15°C until enough young adults were available.

6. Young adults were harvested by washing the culture plates with 1X M9 buffer and collected in a 15 ml centrifuge tube.

7. Worms were washed twice with 1X M9 buffer by centrifuging (using a swinging bucket rotor) the Falcon tube at 1800 rpm for 1 minute to remove any attached bacteria.

8. To the worm pellet, 5 ml of 5% sodium hypochlorite bleach solution was added and incubated for 5-7 minutes with brief vortexing every 2 minutes to break the worm bodies.

9. After 5-7 minutes of incubation, the Falcon tube was centrifuged at 1800 rpm for 1 minute to pellet eggs released from the bodies.

10. The supernatant was removed, and the egg pellet was washed thrice with 10 ml of 1X M9 buffer, each time followed by centrifugation (using a swinging bucket rotor) at 1800 rpm for 1 minute to completely remove the bleach solution.

11. Finally, the egg pellet was resuspended in 12 ml of 1X M9 buffer and incubated for 16 hours at 15°C in a rotor at a speed of 20 rpm to obtain a synchronized L1 larval population.

**Day 1**

**Preparation of NGM RNAi feed**

12. After 16 hours (step 4) of incubation, 4mM IPTG (final concentration) was added to each well for RNAi induction, using a multichannel pipette, and incubated for an additional 1 hour at 37°C, 240 rpm.

13. Bacterial cells were pelleted by centrifuging at 4000 rpm for 10 minutes in a swinging bucket rotor attached with a 96-well adaptor.

14. The supernatant was discarded by rapid inversion to avoid cross-contamination.

15. The bacterial cells were resuspended in 250 µl of NGM medium (half the concentration of the LB used) supplemented with ampicillin (100 µg/ml) and 4 mM IPTG.

16. 60 µl of the NGM RNAi feed was dispensed using a multichannel pipette into each well of flat-bottomed 96-well plates (in triplicates) as feed for the worms.

**RNAi feeding of synchronized L1 larvae**

17. The synchronized population of L1 larval worms obtained after 16 hours (step 11) was diluted to a concentration of 15-20 worms/10 µl drop of 1X M9 by placing worms on slides and counting under a microscope.

18. About 15-20 L1 worms were added to each well of the 96-well flat-bottomed plates containing 60 µl of NGM RNAi feed. Tubes were continuously inverted to prevent any settling of larvae in the 15-ml Falcon tube.

19. The edges of the 96-well plate were sealed with parafilm wrap to prevent the evaporation of feed during incubation.

20. The 96-well plate containing *daf-2(e1370)* L1 larvae was allowed to grow at a sub-permissive temperature of 22.5°C in an incubator shaker at 220 rpm. Note: A few tissue papers soaked in autoclaved water were placed to maintain humidity and prevent evaporation of RNAi feed from the plates.

**Day 4**

**Scoring Dauers**

21. On the day of scoring, the 96-well plates were removed from the incubator, and the lid was removed to wipe any water droplets condensed on the inner surface of the lid for easy visualization of the worms under a stereomicroscope. The lid was then placed back and dauers were scored.

22. Dauers were identified within the liquid culture as dark, thin, and tapered on the edges, aligned straight or in a comma shape in the liquid culture.

23. First, the percentage of dauer formation in each well containing control, *pdk-1*, and *daf-16* RNAi was scored. The entire plate was scored after ensuring controls had worked i.e., *pdk-1* (>90%), and *daf-16* (0%) RNAi. Percentage dauer formation was calculated by dividing the total number of dauers with the total number of worms in a well.

24. Graphs were plotted, with the percentage of dauer formation representing the average of at least two biological repeats.

### Lock-in 96-well lifespan assay

To determine the effect of the positive hits from the dauer-based kinome-wide RNAi screening on lifespan extension, we performed a locked-in 96-well lifespan assay. The protocol is a modification of a previously published high-throughput lifespan screening protocol^40^. We used a synchronized population (obtained as described in step 5 above) of sterile day-1 TJ1060 strains (grown at 25°C) for lifespan analysis to avoid transfers to fresh plates due to progeny production. The 96-well lifespan assay plates were prepared by utilizing an automated 8-channel dispenser. We dispensed 0.15 mL of molten NGM agar (2mM IPTG) into each well of a 96-well plate in a laminar flow cabinet and let the plates dry overnight. Then, we seeded them with 5 mL of 5x concentrated HT115 RNAi bacteria using the automated 8-channel dispenser, as before. The plates were allowed to dry with the lid open for 3-4 hours. To these plates, a synchronized population of day-1 adult TJ1060 worms was added using an automated 8-channel dispenser. The density of the worms in 1x M9 was prepared such that approximately 10-12 worms were dispensed in a 10 µL drop. Plates were allowed to dry for approximately 45 minutes, then covered with lids, and placed in a box at 25°C. Plates were monitored once a week for the first two weeks and then daily thereafter. When the empty vector control had reached > 90% mortality, we scored all the wells for the dead and alive population. Provoked movement by continuously tapping and dropping the plate from a 7 cm height was used to differentiate live worms from dead ones. The clone was recorded as a hit if the percent mortality was <90% and it was reproducible across at least 6 trials out of a total of 10 trial plates. The hits identified from the screen were retested in a 35 mm NGM RNAi plates.

### Lifespan assay in NGM RNAi plates

A synchronized population of worms was obtained from a 2-hour egg lay of day-1 adult hermaphrodites of TJ1060 grown at 15°C. Eggs were then transferred to 25°C and allowed to grow into sterile young adults. On day 1 of adulthood, 50 worms were transferred to either empty vector control or RNAi-seeded 35-mm NGM plates with HT115 bacteria. From day 1, worms were transferred to fresh RNAi-seeded plates every other day until day 7, and then twice over the next 2 weeks. Worms were scored as dead or alive every other day. Worms that failed to display touch-provoked movement were scored as dead. Worms that died from causes other than aging, such as sticking to the plate walls, internal hatching of eggs (’bagging’), or gonadal extrusion, were censored. All lifespan assays were performed at 25°C. Experiments were done in two biological replicates. The worms were scored independently by two investigators who were blinded to the study design and RNAi clone identity. Survival was plotted using GraphPad Prism and p-values determined using Mantel-Cox Log rank test in OASIS: online application for the survival analysis^58^.

### Thermotolerance assay

To determine the effect of each kinase knockdown on thermal stress resistance, we exposed TJ1060 worms to RNAi targeting that kinase at day-1 of adulthood. For each of the two biological replicates, we used two 35 mm NGM RNAi plates, each containing approximately 40 worms. Worms were transferred to fresh RNAi plates every other day until they reached day-7 of adulthood. At this point, they were exposed to a temperature of 35°C for 12 hours. After this period, the worms were scored for survival. Plates were scored by two independent investigators who were blinded to the study design and RNAi clone identity. Survival was plotted using GraphPad Prism and p-values determined using an unpaired Student’s t-test.

## Author contributions

Conception: AM; Investigation: MC, AF, and PS; Data analysis: MC and AF; Original manuscript draft and figures: MC; Review: AM; Technical infrastructure and resources: GL and AM.

## Acknowledgments

The authors thank past and present members of the Molecular Aging Laboratory at the National Institute of Immunology (NII). Special thanks go to Ravi Kumar (NII), Sonu Gupta (NII) and Dr. Dipa Bhaumik (Buck Institute) for their help with laboratory resources. *C. elegans* strains used in this work were provided by the *Caenorhabditis* Genetics Center (CGC), which is funded by the National Institutes of Health (NIH) Office of Research Infrastructure Programs (P40OD010440). Figure 1 and 2A created using BioRender.com. M.C. was supported by a Department of Biotechnology research fellowship and a postdoctoral fellowship from the Larry L. Hillblom Foundation. This work was supported by Ramalingaswami Re-entry Fellowship, National Bioscience Award for Career Development (BT/HRD/NBA/38/04/2016), SERB-STAR award (STR/2019/000064), J.C. Bose National Fellowship (JCB/2022/000021) to A.M.

## Data availability

All data supporting the findings of this study are available in the main text, in the Supplementary Materials. Any unique materials used in the study are available from the authors or from commercially available sources. This study did not use or generate any data code.

## Supplementary Figures and Tables legend

**Supplementary Table 1:** Percentage of *daf-2(e1370)* dauer formation in each well after treatment with RNAi clones targeting 229 kinases. Related to Figure 2B

| RNAi | Percent dauer formation |  |  |  |  |  |
| --- | --- | --- | --- | --- | --- | --- |
| C09B8.7 | 4.76190476 | 4 | 4.34782609 | 13.6363636 | 14.2857143 | 10 |
| Y111B2A.b | 4.54545455 | 8.69565217 | 13.0434783 |  |  |  |
| M79.1 | 4.76190476 | 8.69565217 | 5.55555556 | 3.22580645 | 10.5263158 | 21.7391304 |
| F33D11.7 | 12.5 | 0 | 0 | 25 | 14.2857143 | 8 |
| F22D6.5 |  |  |  | 11.1111111 |  |  |
| T14E8.1 | 5.88235294 | 0 | 7.69230769 | 15.7894737 | 12.5 | 28.5714286 |
| F38E9.5 | 0 | 15 | 11.5384615 | 8.69565217 | 17.8571429 | 21.0526316 |
| ZK632.3 | 4.34782609 | 14.2857143 | 4.54545455 | 16.2790698 | 21.7391304 | 16.6666667 |
| EEED8.9 | 37.5 | 0 | 21.4285714 | 0 | 20 | 0 |
| C36B7.1 | 19.2307692 | 8 | 4.34782609 | 14.2857143 | 23.5294118 | 12.5 |
| C08H9.5 | 0 | 10 | 9.52380952 | 15 | 24 | 26.3157895 |
| B0205.7 | 20 | 6.66666667 | 5.88235294 | 12.5 | 23.0769231 | 18.75 |
| C24G7.5 | 15.7894737 | 16.6666667 | 0 | 24.3902439 | 17.6470588 | 16.1290323 |
| K08F8.1 | 14.8148148 | 7.14285714 | 10.5263158 | 10.7142857 | 20 | 27.7777778 |
| C16D9.2 |  |  |  | 25 | 10 | 11.1111111 |
| C04A11.3 | 21.0526316 | 4.76190476 | 9.09090909 | 9.09090909 | 26.6666667 | 21.875 |
| W03F8.2 | 0 | 12.5 | 11.7647059 | 20 | 26.6666667 | 22.2222222 |
| T11F8.4 | 0 | 6.25 | 5.55555556 | 16.6666667 | 33.3333333 | 33.3333333 |
| F17E5.1 | 25 | 27.2727273 | 14.2857143 | 5.88235294 | 7.14285714 | 17.3913043 |
| C14A4.13 | 6.25 | 10.5263158 | 17.6470588 | 10.5263158 | 26.9230769 | 30.7692308 |
| C39H7.1 | 6.25 | 10.5263158 | 5 | 23.0769231 | 24.2424242 | 35.483871 |
| C05H8.1 | 0 | 11.7647059 | 9.52380952 | 28.125 | 26.6666667 | 30 |
| C08H9.8 | 14.2857143 | 7.14285714 | 5 | 28 | 20 | 31.8181818 |
| F59F5.3 | 0 | 28.5714286 |  | 25.8064516 | 13.3333333 | 20.8333333 |
| K08H2.5 | 7.69230769 | 25 | 13.3333333 | 28.5714286 | 5.55555556 | 30.4347826 |
| D2045.5 | 16.6666667 | 32.2580645 |  | 16.6666667 | 10 | 16.6666667 |
| M03C11.1 | 5.55555556 | 10 | 31.25 | 32.1428571 | 18.75 | 13.5135135 |
| F35G12.3 | 18.1818182 | 11.7647059 | 13.0434783 | 17.8571429 | 29.1666667 | 25 |
| Y69E1A.3 | 9.52380952 | 13.3333333 | 9.375 | 17.6470588 | 22.8571429 | 43.75 |
| ZC504.3 | 22.5 | 5.88235294 | 22.2222222 | 14.2857143 | 20 | 33.3333333 |
| W09C5.5 | 10 | 16.6666667 | 29.4117647 | 17.6470588 | 33.3333333 | 15.3846154 |
| C29H12.5 | 46.6666667 | 25 | 35.7142857 | 0 | 9.09090909 | 6.25 |
| R03G5.2 | 28 | 10 | 19.2307692 | 27.2727273 | 16.6666667 | 21.7391304 |
| W03G1.6 | 7.14285714 | 42.8571429 | 22.2222222 | 16 | 21.4285714 | 13.3333333 |
| B0218.3 | 22.2222222 | 0 | 8.33333333 | 22.7272727 | 31.5789474 | 38.8888889 |
| F59A6.4 | 31.5789474 | 12.5 | 14.8148148 | 18.5185185 | 8.57142857 | 40.625 |
| C01G6.8 | 16.6666667 | 13.3333333 | 26.3157895 | 5.55555556 | 32 | 34.4827586 |
| R06A10.4 | 32.3529412 | 35 | 23.0769231 | 18.1818182 | 8 | 12 |
| F35H8.7 | 8.33333333 | 12 | 10 | 20.5882353 | 44.4444444 | 33.3333333 |
| F36H12.8 | 0 | 12.5 | 5.55555556 | 75 | 17.3913043 | 19.047619 |
| ZK354.2 | 14.2857143 | 12.5 | 13.0434783 | 21.7391304 | 40 | 30.7692308 |
| ZC416.4 | 9.52380952 | 6.4516129 | 11.7647059 | 45.8333333 | 28.125 | 30.7692308 |
| F45H7.4 | 26.6666667 | 18.5185185 | 25 | 22.7272727 | 18.1818182 | 22.7272727 |
| DC2.7 | 5.26315789 | 4.34782609 | 16.6666667 | 60 | 20 | 27.7777778 |
| Y81G3A.3 | 22.7272727 | 15 | 28.5714286 | 30.4347826 | 11.1111111 | 26.3157895 |
| B0478.1 | 7.69230769 | 19.2307692 | 15 | 30.4347826 | 39.1304348 | 23.5294118 |
| R05H5.4 | 19.2307692 | 21.4285714 | 9.09090909 | 45.4545455 | 36 | 6.25 |
| ZC504.4 | 17.3913043 | 29.4117647 | 14.2857143 | 23.8095238 | 28.5714286 | 24 |
| K06H7.8 | 17.6470588 | 14.8148148 | 3.38983051 | 43.3333333 | 29.0322581 | 29.4117647 |
| E02H4.6 | 17.6470588 | 15.3846154 | 10 | 47.3684211 | 29.6296296 | 18.75 |
| C05D2.1 | 40 | 11.7647059 | 31.8181818 | 10 | 22.7272727 | 22.7272727 |
| C44H4.6 | 18.1818182 | 15.3846154 | 20 | 22.2222222 | 32.1428571 | 33.3333333 |
| ZC581.1 |  |  |  | 17.6470588 | 36.3636364 | 16.6666667 |
| R02C2.1 | 6.06060606 | 3.57142857 | 14.2857143 | 45.8333333 | 26.3157895 | 45.7142857 |
| R13H9.6 | 33.3333333 | 5.55555556 | 7.69230769 | 33.3333333 | 15.3846154 | 47.3684211 |
| F36H12.9 | 12.5 | 8.33333333 | 10.5263158 | 31.5789474 | 42.8571429 | 37.5 |
| C34B2.3 | 33.3333333 | 30 | 44.4444444 | 0 | 23.0769231 | 12.5 |
| K11H12.9 | 8.33333333 | 0 | 20 | 38.0952381 | 42.8571429 | 34.7826087 |
| K07A9.2 | 4.34782609 | 0 | 5 | 34.7826087 | 50 | 50 |
| C09B9.4 | 21.0526316 | 28.5714286 | 36.8421053 | 15 | 15 | 18.1818182 |
| T27E9.3 | 0 | 8.69565217 | 11.1111111 | 22.5806452 | 47.0588235 | 55.5555556 |
| F46G11.3 | 12 | 17.3913043 | 6.66666667 | 53.3333333 | 25 | 30.7692308 |
| F13B9.4 | 12.9032258 | 20.6896552 | 26.0869565 | 25 | 24 | 36.8421053 |
| T01H10.4 | 18.1818182 | 11.5384615 | 15.7894737 | 31.8181818 | 33.3333333 | 35 |
| F26A1.4 | 55.5555556 | 28.5714286 | 26.6666667 | 5.88235294 | 6.66666667 | 23.3333333 |
| F22H10.5 | 23.8095238 | 0 | 0 | 27.2727273 | 43.75 | 51.8518519 |
| F46F6.2 | 20.8333333 | 41.9354839 | 23.3333333 | 13.3333333 | 15.3846154 | 33.3333333 |
| F53G12.6 | 25 | 40 | 46.1538462 | 27.2727273 | 10 | 0 |
| Y119D3_455.b | 29.16667 | 57.14286 | 37.5 | 5 | 16.6666667 | 5 |
| T05G5.3 | 26.6666667 | 10.5263158 | 15.7894737 | 39.2857143 | 35.2941176 | 23.3333333 |
| M176.7 | 20.8333333 | 9.52380952 | 9.09090909 | 40 | 18.75 | 53.3333333 |
| C24A1.3 | 13.0434783 | 0 | 9.09090909 | 36.3636364 | 32 | 61.1111111 |
| F43C1.2 | 28.57143 | 22.22222 | 25 | 30 | 14.2857143 | 31.8181818 |
| R13H9.5 | 40 | 18.75 | 30.7692308 | 17.8571429 | 33.3333333 | 12 |
| C07A9.3 | 30 | 25 | 22.7272727 | 15.3846154 | 28.5714286 | 22.7272727 |
| C29F9.7 | 11.7647059 | 10.5263158 | 5 | 40.7407407 | 52.173913 | 34.6153846 |
| R03D7.5 | 9.52380952 | 4.34782609 | 13.3333333 | 31.25 | 42.8571429 | 54.5454545 |
| R13F6.7 | 7.69230769 | 9.09090909 | 9.52380952 | 50 | 35 | 50 |
| F31E3.2 | 14.2857143 | 19.2307692 | 20 | 50 | 35 | 40 |
| R09D1.12 | 21.0526316 | 10.5263158 | 15.7894737 | 28.5714286 | 47.826087 | 40 |
| F57B9.8 | 6.25 | 15 | 0 | 10 | 10 | 17.3913043 |
| F25F2.1 | 33.3333333 | 31.0344828 | 26.6666667 | 15.7894737 | 26.3157895 | 23.5294118 |
| C03C10.1 | 16.6666667 | 16.6666667 | 11.7647059 | 16.6666667 | 36.8421053 | 66.6666667 |
| ZC449.3 | 13.0434783 | 25 | 13.7931034 | 18.75 | 36.3636364 | 59.2592593 |
| F35C8.3 | 9.52380952 | 0 | 15.3846154 | 76.9230769 | 19.047619 | 45.4545455 |
| T05A7.6 | 4 | 6.66666667 | 9.09090909 | 0 | 27.2727273 | 21.0526316 |
| B0545.1a | 29.4117647 | 41.6666667 | 23.0769231 | 33.3333333 | 17.1428571 | 22.7272727 |
| C25F6.4 | 20 | 14.8148148 | 11.7647059 | 55.2631579 | 28.125 | 38.0952381 |
| ZK1067.1 | 11.1111111 | 13.3333333 | 11.7647059 | 50 | 40.9090909 | 41.1764706 |
| W02B12.12 | 14.2857143 | 26.3157895 | 16.6666667 | 4.16666667 | 61.1111111 | 47.0588235 |
| B0464.5 | 6.25 | 24.3243243 | 22.2222222 | 33.3333333 | 30.4347826 | 55 |
| K04C1.5 | 0 | 21.0526316 | 0 | 41.9354839 | 37.5 | 72.2222222 |
| M176.6 | 13.6363636 | 35.2941176 | 18.75 | 33.3333333 | 47.826087 | 24.137931 |
| F19H6.1 | 5 | 0 | 16 | 9.67741935 | 3.125 | 8.57142857 |
| C28A5.6 | 50 | 30 | 33.3333333 | 25 | 12.5 | 22.7272727 |
| C05D10.2 | 57.14286 | 26.66667 | 27.77778 | 27.7777778 | 15.3846154 | 19.2307692 |
| F19C6.1 | 30.2325581 | 15.7894737 | 18.9189189 | 36.3636364 | 35.2941176 | 37.5 |
| C06E8.3 | 30 | 15.7894737 | 19.047619 | 40 | 32 | 38.0952381 |
| W10G6.2 | 0 | 20 | 10 | 30 | 65.5172414 | 50 |
| F59F3.1 | 14.8148148 | 25 | 27.2727273 | 41.025641 | 41.6666667 | 26.0869565 |
| C18H7.4 | 11.1111111 | 10.5263158 | 25.9259259 | 40 | 46.6666667 | 42.3076923 |
| Y39G8C.2 | 33.3333333 | 13.3333333 | 50 | 25 | 37.5 | 18.75 |
| F59E12.2 | 50 | 45.8333333 | 57.1428571 | 12.5 | 13.3333333 | 0 |
| F58A3.2 | 33.3333333 | 7.14285714 | 18.4210526 | 21.875 | 48.1481481 | 50 |
| C55C3.4 | 19.047619 | 18.75 | 14.2857143 | 36.3636364 | 46.1538462 | 44.4444444 |
| T05E8.2 | 31.25 | 44.8275862 | 59.2592593 | 16.6666667 | 13.3333333 | 14.2857143 |
| R107.4 | 27.7777778 | 33.3333333 | 42.8571429 | 35.483871 | 16.6666667 | 23.8095238 |
| F46C3.1 | 5.88235294 | 8.69565217 | 24.2424242 | 53.8461538 | 44.4444444 | 42.8571429 |
| H25P06.2a | 60.8695652 | 55 | 50 | 0 | 0 | 14.2857143 |
| C41C4.4 | 33.3333333 | 68.75 | 54.5454545 | 13.3333333 | 5.26315789 | 5.55555556 |
| F54F7.5 | 20 | 13.6363636 | 14.8148148 | 50 | 48 | 35 |
| C03C10.2 | 20 | 13.0434783 | 29.1666667 | 44.4444444 | 42.8571429 | 32.1428571 |
| F49C5.4 | 22.22222 | 8.333333 | 29.16667 | 41.6666667 | 26.9230769 | 53.8461538 |
| ZK596.2 | 32.3529412 | 32.2580645 | 43.75 | 22.2222222 | 31.25 | 20.8333333 |
| F42A10.4 | 25 | 25 | 42.1052632 | 38.8888889 | 31.25 | 21.4285714 |
| Y4C6A.k | 20.8333333 | 10.5263158 | 46.6666667 | 50 | 43.4782609 | 13.7931034 |
| ZK930.1 | 6.66666667 | 13.3333333 | 10.5263158 | 67.7419355 | 13.0434783 | 42.3076923 |
| F28B12.3 | 41.9354839 | 35.2941176 | 30 | 25 | 36.8421053 | 18.1818182 |
| R11E3.1 |  |  |  |  |  |  |
| D2024.1 | 15.625 | 18.75 | 10 | 55.8823529 | 53.8461538 | 35.2941176 |
| F55A8.2 | 20 | 4.16666667 |  | 43.4782609 | 43.75 | 47.0588235 |
| F15A2.6 | 33.3333333 | 23.8095238 |  | 41.3793103 | 34.7826087 | 25.9259259 |
| M176.9 | 5 | 30 | 24 | 41.6666667 | 50 | 59.2592593 |
| F35C8.2 | 7.69230769 |  | 7.40740741 | 55.5555556 | 38.7096774 | 51.6129032 |
| C24G6.2 | 23.0769231 | 6.66666667 | 9.67741935 | 41.6666667 | 55 | 57.1428571 |
| C35E7.10 | 20 | 0 | 25 | 55.5555556 | 34.7826087 | 61.1111111 |
| T25E12.4 | 15.3846154 | 7.40740741 | 14.2857143 | 44 | 58.8235294 | 57.1428571 |
| M04C9.5 | 62.962963 | 51.8518519 | 42.8571429 | 20 | 0 | 21.4285714 |
| T04C10.1 | 10 | 20 | 31.5789474 | 63.1578947 | 25 | 50 |
| W02B3.2 | 31.5789474 | 21.4285714 | 25 |  | 38.8888889 | 50 |
| K08A8.1 | 23.5294118 | 25 | 53.3333333 | 26.3157895 | 33.3333333 | 39.2857143 |
| E02H4.3 | 22.2222222 | 36 | 19.047619 | 50 | 27.7777778 | 48.1481481 |
| ZK354.6 | 27.2727273 | 16.6666667 | 25 | 44.8275862 | 40.625 | 50 |
| Y54E10B_152. | 75 | 51.3513514 | 44.4444444 | 11.7647059 | 5.88235294 | 17.6470588 |
| W03A5.1 | 11.1111111 | 5.26315789 | 12.5 | 65 | 65.2173913 | 47.0588235 |
| F40A3.5 | 41.1764706 | 10.7142857 | 26.9230769 | 46.4285714 | 40.5405405 | 40.7407407 |
| F42G10.2 | 40.9090909 | 35 | 11.5384615 | 56.25 | 28.5714286 | 34.2857143 |
| F49E11.1 |  |  |  | 65.5172414 | 9.09090909 | 29.4117647 |
| K08B4.5 | 25 | 26.3157895 | 8.33333333 | 31.5789474 | 51.7241379 | 65.2173913 |
| F22H10.1 | 12.5 | 18.1818182 |  | 40.9090909 | 18.1818182 | 26.0869565 |
| F59F3.5 | 10.5263158 | 23.8095238 | 29.6296296 | 52.9411765 | 38.0952381 | 57.6923077 |
| T01C8.1 | 22.2222222 | 33.3333333 | 23.0769231 | 43.75 | 37.5 | 52.9411765 |
| ZK938.5 | 7.14285714 | 18.75 | 26.6666667 | 15.3846154 | 30.4347826 | 50 |
| C44C10.7 | 42.1052632 | 26.0869565 | 52.173913 | 27.2727273 | 27.5862069 | 39.2857143 |
| K11E8.1 | 23.3333333 | 10.7142857 | 22.2222222 | 27.2727273 | 52.173913 | 44.4444444 |
| Y52D3.1 | 23.8095238 | 33.3333333 | 22.2222222 | 52.173913 | 50 | 10 |
| Y51B9A.9 | 8.33333333 | 6.4516129 | 18.1818182 | 33.3333333 | 22.2222222 | 15 |
| T27C10.5 | 41.3793103 | 67.5675676 | 59.2592593 | 25 | 14.2857143 | 15 |
| D2045.7 | 44.4444444 | 8.33333333 | 30.4347826 | 71.875 | 34.6153846 | 33.3333333 |
| F09A5.2 | 15 | 12.5 | 12.5 | 13.3333333 | 15.3846154 | 35.7142857 |
| C34F11.5 | 51.7241379 | 71.4285714 | 57.1428571 | 9.09090909 | 22.2222222 | 13.3333333 |
| T07A9.3 | 23.8095238 | 22.7272727 | 14.2857143 | 30.4347826 | 50 | 85.7142857 |
| Y39G8B.f | 30.7692308 | 35.7142857 | 26.6666667 | 58.0645161 | 40 | 36.3636364 |
| PAR2.3 | 4.76190476 | 27.7777778 | 17.6470588 | 52.173913 | 40 | 54.8387097 |
| VZC374L.1 | 22.2222222 | 14.8148148 | 9.52380952 | 64.7058824 | 52.9411765 | 64.2857143 |
| B0207.4 | 38.0952381 | 30 | 34.7826087 | 37.1428571 | 60.6060606 | 28 |
| F08F1.1 | 10.5263158 | 8 | 4.34782609 | 43.75 | 30 | 27.5862069 |
| F16B12.5 | 61.9047619 | 50 | 64.7058824 | 51.8518519 | 43.75 | 42.3076923 |
| H05L14.1 | 40 | 62.5 | 47.3684211 | 33.3333333 | 39.1304348 | 56.25 |
| K02E10.7 | 60.8695652 | 55 | 50 | 38.8888889 | 17.6470588 | 11.7647059 |
| F18H3.5 | 10.5263158 | 31.5789474 | 37.5 | 54.8387097 | 52.173913 | 47.826087 |
| Y38H8A.3 | 75 | 51.3513514 | 44.4444444 | 30 | 15.7894737 | 20 |
| F35C8.1 | 27.7777778 | 38.8888889 | 6.66666667 | 60 | 59.0909091 | 44.4444444 |
| F40G9.13 | 15.7894737 | 21.7391304 | 37.5 | 53.8461538 | 60 | 50 |
| ZK370.5 | 18.75 | 21.0526316 | 20 | 59.2592593 | 46.1538462 | 73.6842105 |
| F26A1.3 | 9.09090909 | 7.69230769 | 9.52380952 | 68.1818182 | 46.1538462 | 54.5454545 |
| ZC581.9 | 63.8888889 | 76.9230769 | 63.1578947 | 25 | 0 | 11.1111111 |
| F09E5.1 | 72.7272727 | 60 | 69.2307692 | 21.0526316 | 7.14285714 | 12.5 |
| T17A3.8 | 10 | 5.55555556 | 0 | 33.3333333 | 35 | 52 |
| C44C8.6 | 25 | 26.9230769 | 35 | 30.4347826 | 19.2307692 | 45 |
| B0523.1 | 41.66667 | 44.44444 | 44.44444 | 31.25 | 29.4117647 | 25 |
| T23G7.5 | 36.3636364 | 29.4117647 | 36.8421053 | 55 | 42.1052632 | 45.1612903 |
| T17E9.1 | 60 | 52.9411765 | 42.1052632 | 32.1428571 | 38.0952381 | 52.173913 |
| C35D10.4 | 11.7647059 | 12 | 5.55555556 | 30 | 40 | 80 |
| F29C4.1 | 11.7647059 | 28.5714286 | 28.5714286 | 55.2631579 | 56.5217391 | 60.8695652 |
| W04G5.6 | 33.3333333 | 6.66666667 | 12.5 | 66.6666667 | 77.7777778 | 57.8947368 |
| Y106G6A.1 | 66.6666667 | 44.4444444 | 54.5454545 | 42.8571429 | 19.047619 | 30 |
| C09G4.2 | 63.8888889 | 76.9230769 | 63.1578947 | 27.2727273 | 10 | 16.6666667 |
| H37N21.1 | 75 | 50 | 40 | 30 | 37.5 | 27.2727273 |
| C56C10.6 | 75 | 84.2105263 | 40 | 12.5 | 25 | 23.0769231 |
| T06C10.3 | 28.5714286 | 73.6842105 | 53.8461538 | 27.7777778 | 16.6666667 | 60 |
| F32D8.1 | 63.3333333 | 37.9310345 | 43.4782609 | 36.6666667 | 28.125 | 51.4285714 |
| Y53C12A.1 | 52.9411765 | 50 | 52.6315789 | 39.1304348 | 27.2727273 | 39.3939394 |
| Y42A5A.4 | 30 | 41.1764706 | 57.1428571 | 46.6666667 | 36.8421053 | 50 |
| D1073.1 | 47.0588235 | 46.1538462 | 38.0952381 | 64 | 35.7142857 | 36.1111111 |
| B0496.3 | 16.6666667 | 44.8275862 | 38.4615385 | 65 | 53.3333333 | 50 |
| C01H6.9 | 64.2857143 | 66.6666667 | 92 | 10 | 23.0769231 | 13.3333333 |
| D1044.3 | 16.6666667 | 0 | 0 | 47.826087 | 68.4210526 | 70 |
| C45G9.1 | 16.6666667 | 3.84615385 | 7.69230769 | 70 | 73.3333333 | 60 |
| W06F12.3 | 13.3333333 | 11.7647059 | 9.09090909 | 36.3636364 | 62.962963 | 54.5454545 |
| R151.4 | 50 | 42.1052632 | 58.8235294 | 47.0588235 | 18.1818182 | 56.7567568 |
| ZC373.3 | 55.5555556 | 33.3333333 | 46.1538462 | 36.3636364 | 39.1304348 | 64 |
| C50H2.7 | 62.962963 | 51.8518519 | 42.8571429 | 37.5 | 44.4444444 | 36.3636364 |
| W08D2.8 | 41.6666667 | 90 | 77.7777778 | 21.4285714 | 7.14285714 | 38.4615385 |
| T09B4.7 | 28.5714286 | 73.6842105 | 53.8461538 | 37.5 | 41.6666667 | 42.8571429 |
| F22B3.8 | 75 | 50 | 40 | 50 | 30 | 33.3333333 |
| H39E23.1 | 33.3333333 | 30 | 44.4444444 | 43.75 | 81.8181818 | 45.4545455 |
| F39F10.3 | 48.4848485 | 64.2857143 | 5.26315789 | 44.4444444 | 44.4444444 | 75 |
| W06F12.1 | 39.3939394 | 48.5714286 |  | 47.6190476 | 47.6190476 | 52 |
| C54G4.1 | 0 | 8.33333333 | 15.3846154 | 25 | 25 | 31.5789474 |
| W04B5.5 | 17.6470588 | 15.7894737 | 17.3913043 | 26.6666667 | 41.1764706 | 52 |
| F55G1.8 | 21.7391304 | 38.8888889 | 38.0952381 | 61.9047619 | 64.2857143 | 66.6666667 |
| E01H11.1 | 38.8888889 | 21.4285714 | 60.7142857 | 56.5217391 | 63.3333333 | 58.3333333 |
| ZK622.1 | 66.6666667 | 66.6666667 | 90.9090909 | 25 | 23.5294118 | 26.6666667 |
| B0198.3 | 5.55555556 | 21.0526316 | 33.3333333 | 21.0526316 | 12.5 | 26.9230769 |
| T20F10.1 | 57.1428571 | 100 | 95.6521739 | 11.7647059 | 16.6666667 | 22.2222222 |
| C10C6.1 | 64.2857143 | 66.6666667 | 92 | 21.4285714 | 30 | 33.3333333 |
| T01H8.1 | 31.5789474 | 52.173913 | 58.3333333 | 48.1481481 | 66.6666667 | 52.6315789 |
| F22F1.2 | 62.962963 | 54.1666667 |  | 50 | 47.0588235 | 44.4444444 |
| R166.5 | 21.4285714 | 42.8571429 | 5.26315789 | 50 | 66.6666667 | 76 |
| F20B6.8 | 41.3793103 | 67.5675676 | 59.2592593 | 22.2222222 | 63.6363636 | 60 |
| R10D12.10 | 66.6666667 | 44.4444444 | 54.5454545 | 47.6190476 | 18.1818182 | 83.3333333 |
| B0285.1 | 9.52380952 | 11.7647059 | 15.3846154 | 73.3333333 | 86.9565217 | 88.8888889 |
| T22B11.3 | 20 | 26.92308 | 38.09524 | 61.9047619 | 62.0689655 | 57.8947368 |
| R06C7.8 | 41.6666667 | 90 | 77.7777778 | 46.1538462 | 31.25 | 36.3636364 |
| F18F11.5 | 72.2222222 | 88.2352941 | 93.75 | 9.09090909 | 29.1666667 | 47.5 |
| C25A8.5 | 33.3333333 | 68.75 | 54.5454545 | 29.4117647 | 25 | 42.1052632 |
| ZK909.2 | 21.4285714 | 23.3333333 |  | 84.8484848 | 74.0740741 | 72.4137931 |
| T15B12.2 | 10 | 40 | 9.09090909 | 100 | 95.2380952 | 68.1818182 |
| C46C2.1 |  |  |  | 45.4545455 | 66.6666667 |  |
| K06H7.1 | 28.5714286 | 6.25 | 20.8333333 | 95.2380952 | 91.6666667 | 91.6666667 |
| F52C12.2 | 14.2857143 | 5 | 10 | 72.7272727 | 76.1904762 | 78.3783784 |
| C32D5.2 | 95.8333333 | 87.5 | 88.8888889 | 40 | 26.6666667 | 25 |
| F46F2.2 | 56.25 | 55 | 55 | 70 | 76.9230769 | 55.1724138 |
| C03B1.5 | 58.974359 | 51.8518519 | 52 | 90 | 60 | 56.5217391 |
| F28H6.1 | 50 | 50 | 51.8518519 | 66.6666667 | 68.9655172 | 87.0967742 |
| Y106G6E.6 | 90.9090909 | 93.75 | 85.7142857 | 26.6666667 | 42.8571429 | 36.3636364 |
| ZK617.1 | 60.8695652 | 57.6923077 | 54.5454545 | 87.8787879 | 73.0769231 | 85.1851852 |
| K09A9.1 | 100 | 100 | 95 | 100 | 78.2608696 | 85.7142857 |

**Supplementary Table 2:** Percent change in the lifespan relative to empty vector control RNAi. p-value determined using Mantel-Cox log rank test. Strain: TJ1060. Temperature: 25°C. Post-developmental RNAi treatment. Related to Figure 3A and 3B.

| | Mean lifespan in days ± SEM | Median lifespan in days | N | % Mean lifespan change wrt Control | p-value wrt control RNAi |
| --- | --- | --- | --- | --- | --- |
| EV-Control RNAi | 10.27 ± 0.42 | 9.00 | 74 | | |
| *abl-1* | 11.02 ± 0.35 | 12.00 | 90 | 7.3 | 0.2598 |
| *akt-2* | 12.95 ± 0.52 | 12.00 | 80 | 26.1 | 0.0001 |
| *C25A8.5* | 11.21 ± 0.47 | 12.00 | 68 | 9.2 | 0.1792 |
| *C50H2.7* | 10.68 ± 0.36 | 9.00 | 101 | 4.0 | 0.6729 |
| *F32D8.1* | 11.36 ± 0.57 | 12.00 | 42 | 10.6 | 0.2401 |
| *H05L14.1* | 11.37 ± 0.29 | 12.00 | 87 | 10.7 | 0.1389 |
| *kin-1* | 12.05 ± 0.46 | 12.00 | 79 | 17.3 | 0.0066 |
| *kin-23* | 11 ± 0.42 | 12.00 | 64 | 7.1 | 0.3529 |
| *rskn-1* | 11.57 ± 0.37 | 12.00 | 87 | 12.7 | 0.0388 |
| *trk-1* | 11.6 ± 0.54 | 12.00 | 70 | 13.0 | 0.0387 |
| EV-Control RNAi | 10.5 ± 0.28 | 9.00 | 80 | | |
| *T09B4.7* | 11.74 ± 0.39 | 12.00 | 43 | 11.8 | 0.0183 |
| *Y106G6A.1* | 10.8 ± 0.33 | 12.00 | 76 | 2.9 | 0.4593 |
| *ZK622.1* | 11.49 ± 0.32 | 12.00 | 86 | 9.4 | 0.0153 |
| *C56C10.6* | 11.45 ± 0.31 | 12.00 | 75 | 9.0 | 0.0348 |
| *W04B5.5* | 10.61 ± 0.35 | 9.00 | 80 | 1.0 | 0.5413 |
| *T15B12.2* | 13.2 ± 0.43 | 14.00 | 46 | 25.7 | p<0.00001 |
| *F26A1.3* | 11.02 ± 0.32 | 12.00 | 84 | 5.0 | 0.1935 |
| *R151.4* | 10.73 ± 0.32 | 9.00 | 79 | 2.2 | 0.4103 |
| *K06H7.1* | 11.26 ± 0.36 | 12.00 | 80 | 7.2 | 0.0433 |
| *W06F12.3* | 11.66 ± 0.43 | 12.00 | 38 | 11.0 | 0.0379 |
| *F52C12.2* | 11.61 ± 0.30 | 12.00 | 84 | 10.6 | 0.0111 |
| *T22B11.3* | 11.72 ± 0.35 | 12.00 | 85 | 11.6 | 0.0048 |
| *T06C10.3* | 11.52 ± 0.37 | 12.00 | 79 | 9.7 | 0.0171 |
| *F22B3.8* | 10.63 ± 0.32 | 12.00 | 84 | 1.2 | 0.5846 |
| *C03B1.5* | 11.4 ± 0.46 | 12.00 | 45 | 8.6 | 0.0674 |
| *F22F1.2* | 11.15 ± 0.35 | 12.00 | 79 | 6.2 | 0.0848 |
| *F16B12.5* | 10.5 ± 0.32 | 9.00 | 86 | 0.0 | 0.7626 |
| *F09A5.2* | 11.76 ± 0.35 | 12.00 | 85 | 12.0 | 0.0026 |
| *F39F10.3* | 10.8 ± 0.34 | 9.00 | 85 | 2.9 | 0.3600 |
| *ZC373.3* | 10.95 ± 0.30 | 12.00 | 82 | 4.3 | 0.2698 |
| *F33D11.7* | 10.27 ± 0.56 | 9.00 | 41 | -2.2 | 0.9727 |

Strain: TJ1060; Temperature: 25°C; post-development RNAi
The cumulative results of two independent trials, each scored by two investigators who were blinded to clone identity

**Supplementary Figure 1:**
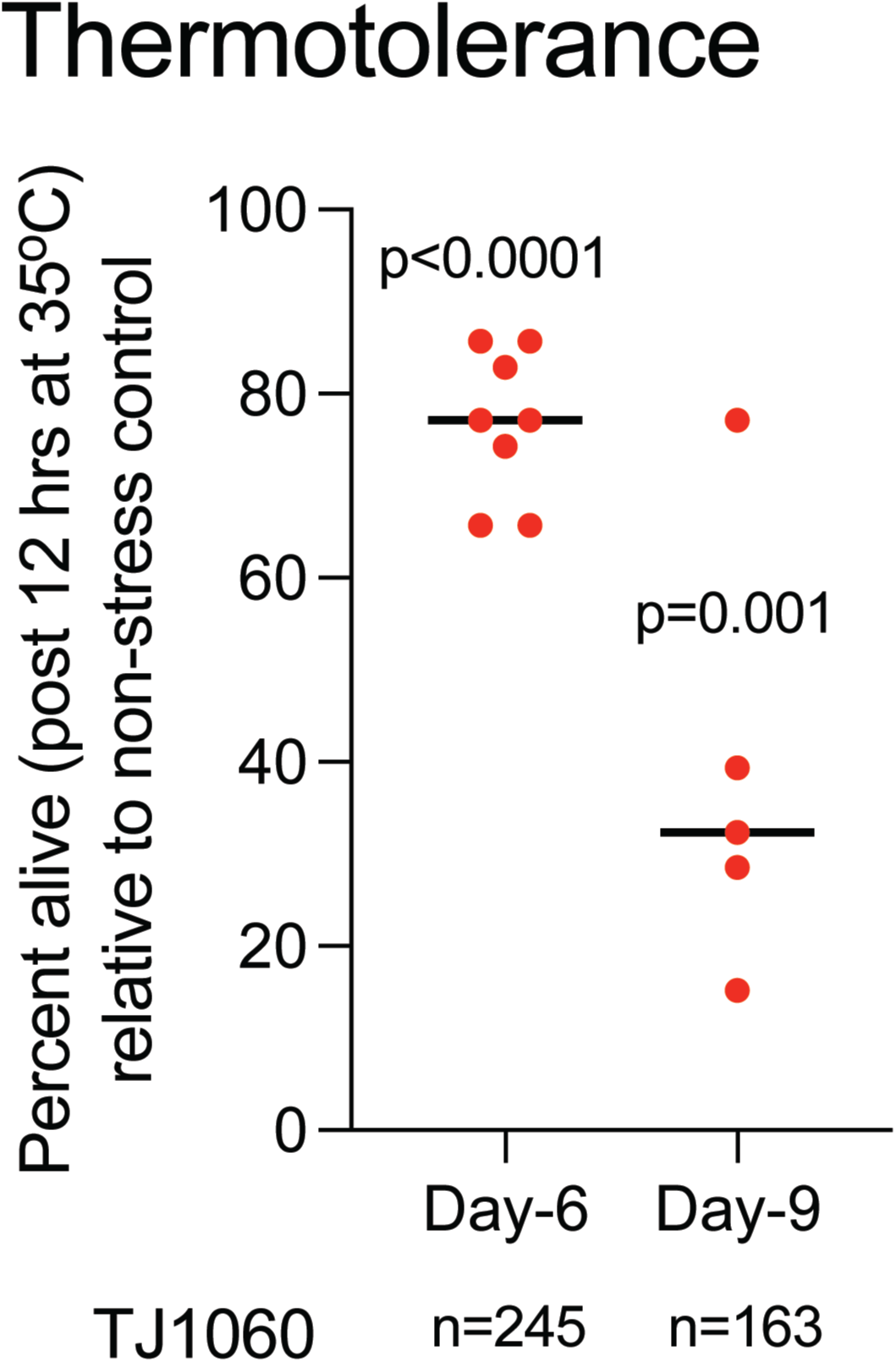
**(A) Thermotolerance assay.** Percent survival of TJ1060 worms after exposure to 35°C for 12 hours. The assay was performed at two different ages: day 6 and day 9 of adulthood. Values are plotted relative to age-matched non-stressed worms. Each dot represents an individual plate with approximately 35 worms across two independent trials. p-values were determined using an unpaired t-test (two-tailed).

**Supplementary Table 3:**
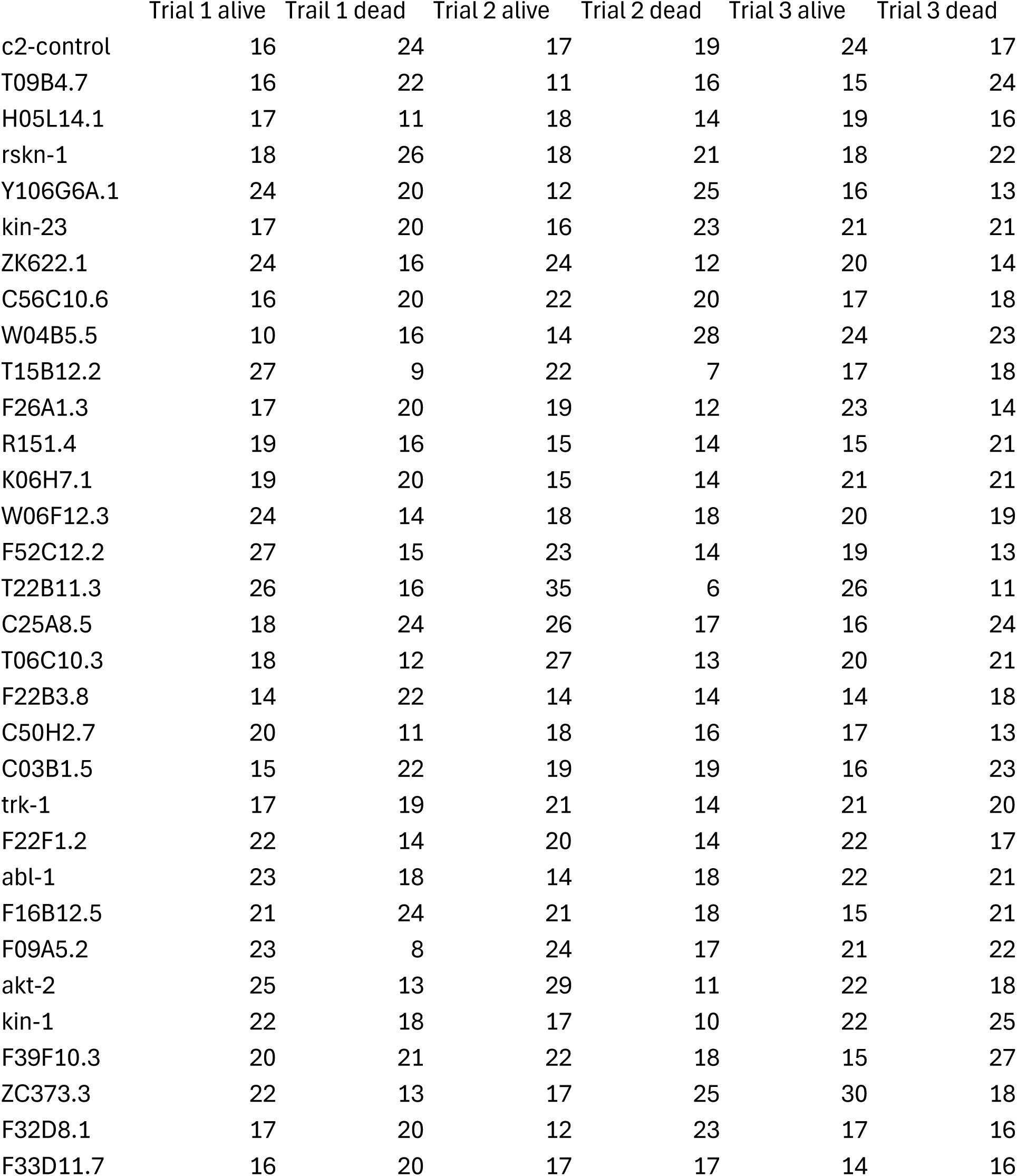
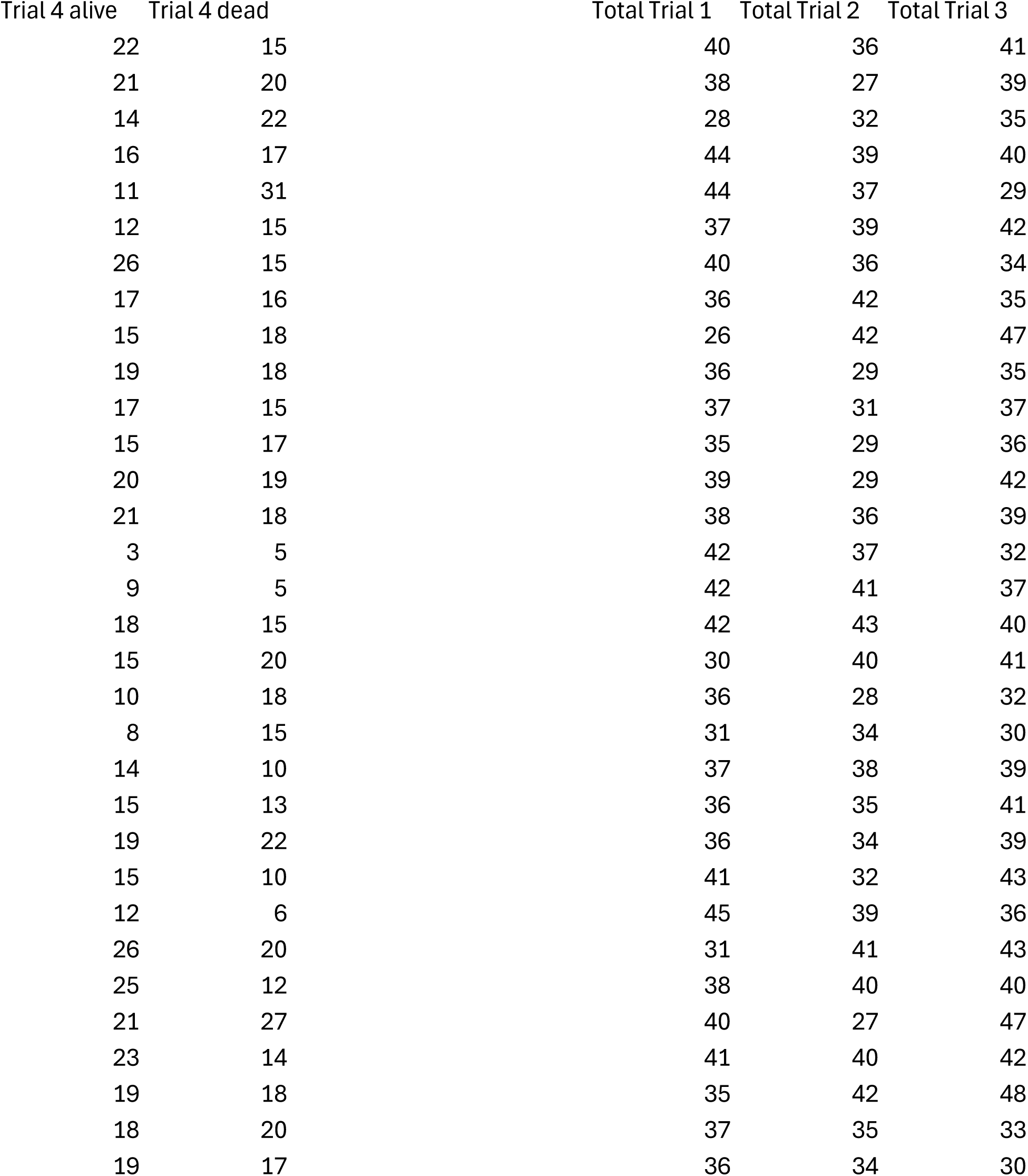
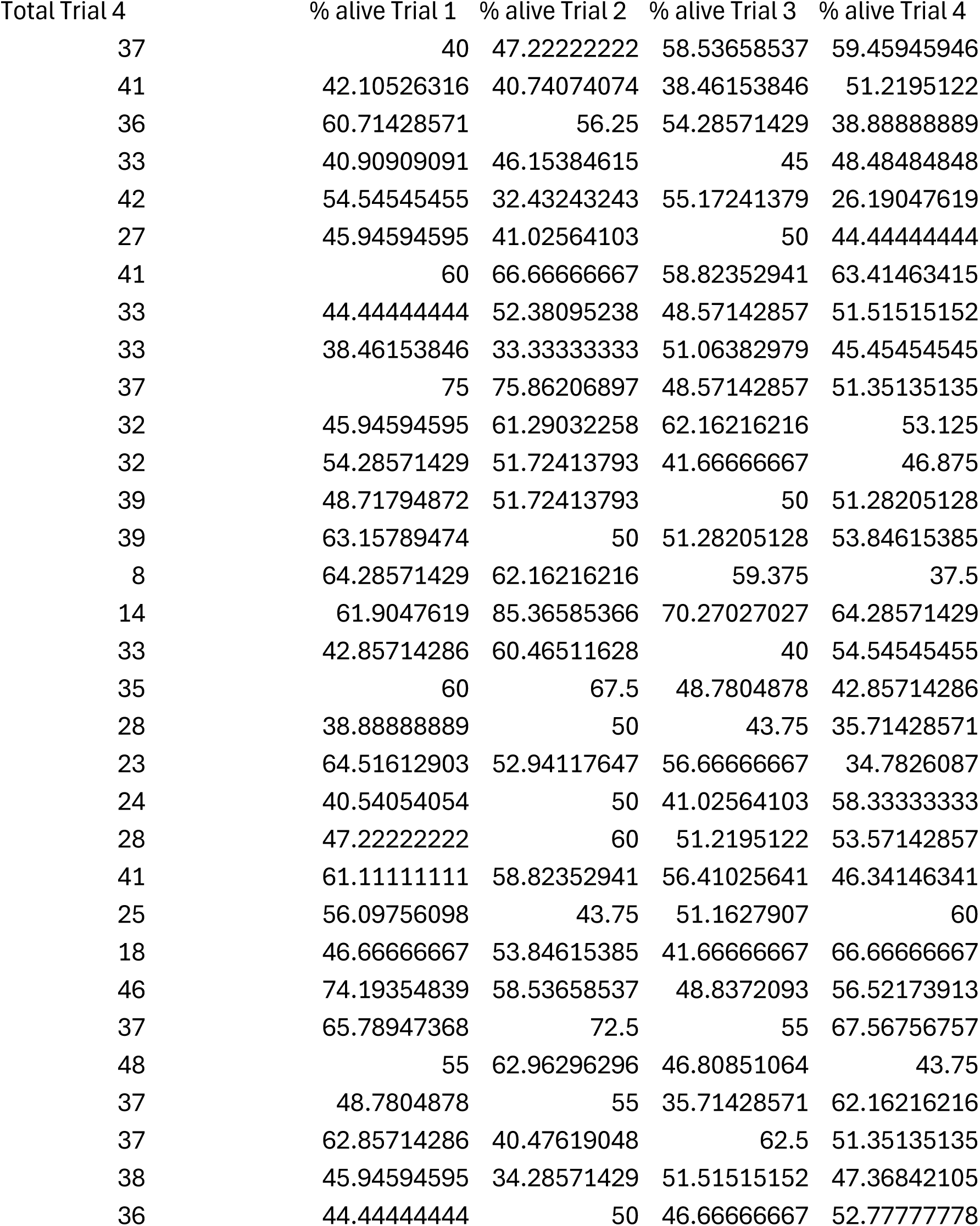
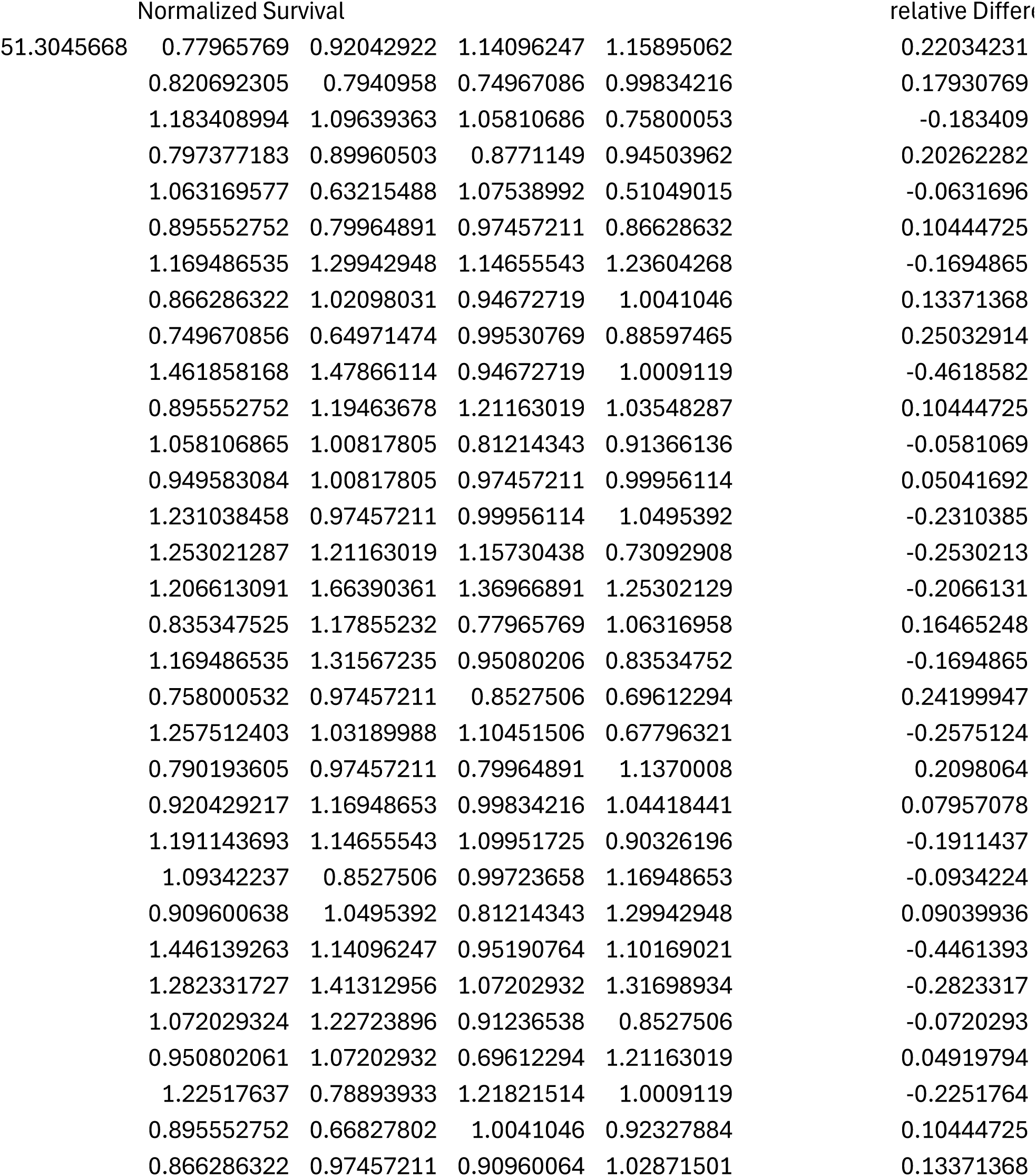
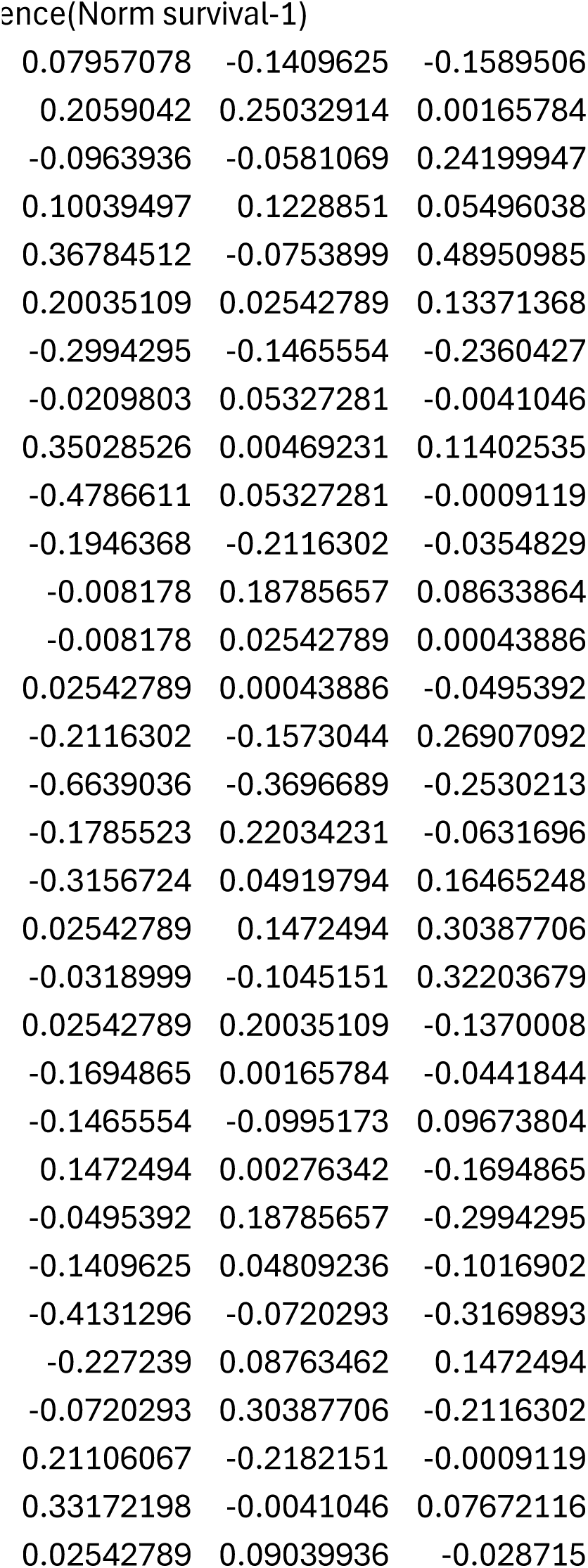
Raw data and calculated percent survival of TJ1060 worms after exposure to 35°C for 12 hours across all trials. Related to Figure 3C.

|  | Trial 1 alive | Trial 1 dead | Trial 2 alive | Trial 2 dead | Trial 3 alive | Trial 3 dead |
| --- | --- | --- | --- | --- | --- | --- |
| c2-control | 16 | 24 | 17 | 19 | 24 | 17 |
| T09B4.7 | 16 | 22 | 11 | 16 | 15 | 24 |
| H05L14.1 | 17 | 11 | 18 | 14 | 19 | 16 |
| rskn-1 | 18 | 26 | 18 | 21 | 18 | 22 |
| Y106G6A.1 | 24 | 20 | 12 | 25 | 16 | 13 |
| kin-23 | 17 | 20 | 16 | 23 | 21 | 21 |
| ZK622.1 | 24 | 16 | 24 | 12 | 20 | 14 |
| C56C10.6 | 16 | 20 | 22 | 20 | 17 | 18 |
| W04B5.5 | 10 | 16 | 14 | 28 | 24 | 23 |
| T15B12.2 | 27 | 9 | 22 | 7 | 17 | 18 |
| F26A1.3 | 17 | 20 | 19 | 12 | 23 | 14 |
| R151.4 | 19 | 16 | 15 | 14 | 15 | 21 |
| K06H7.1 | 19 | 20 | 15 | 14 | 21 | 21 |
| W06F12.3 | 24 | 14 | 18 | 18 | 20 | 19 |
| F52C12.2 | 27 | 15 | 23 | 14 | 19 | 13 |
| T22B11.3 | 26 | 16 | 35 | 6 | 26 | 11 |
| C25A8.5 | 18 | 24 | 26 | 17 | 16 | 24 |
| T06C10.3 | 18 | 12 | 27 | 13 | 20 | 21 |
| F22B3.8 | 14 | 22 | 14 | 14 | 14 | 18 |
| C50H2.7 | 20 | 11 | 18 | 16 | 17 | 13 |
| C03B1.5 | 15 | 22 | 19 | 19 | 16 | 23 |
| trk-1 | 17 | 19 | 21 | 14 | 21 | 20 |
| F22F1.2 | 22 | 14 | 20 | 14 | 22 | 17 |
| abl-1 | 23 | 18 | 14 | 18 | 22 | 21 |
| F16B12.5 | 21 | 24 | 21 | 18 | 15 | 21 |
| F09A5.2 | 23 | 8 | 24 | 17 | 21 | 22 |
| akt-2 | 25 | 13 | 29 | 11 | 22 | 18 |
| kin-1 | 22 | 18 | 17 | 10 | 22 | 25 |
| F39F10.3 | 20 | 21 | 22 | 18 | 15 | 27 |
| ZC373.3 | 22 | 13 | 17 | 25 | 30 | 18 |
| F32D8.1 | 17 | 20 | 12 | 23 | 17 | 16 |
| F33D11.7 | 16 | 20 | 17 | 17 | 14 | 16 |

| Trial 4 alive | Trial 4 dead | Total Trial 1 | Total Trial 2 | Total Trial 3 |
| --- | --- | --- | --- | --- |
| 22 | 15 | 40 | 36 | 41 |
| 21 | 20 | 38 | 27 | 39 |
| 14 | 22 | 28 | 32 | 35 |
| 16 | 17 | 44 | 39 | 40 |
| 11 | 31 | 44 | 37 | 29 |
| 12 | 15 | 37 | 39 | 42 |
| 26 | 15 | 40 | 36 | 34 |
| 17 | 16 | 36 | 42 | 35 |
| 15 | 18 | 26 | 42 | 47 |
| 19 | 18 | 36 | 29 | 35 |
| 17 | 15 | 37 | 31 | 37 |
| 15 | 17 | 35 | 29 | 36 |
| 20 | 19 | 39 | 29 | 42 |
| 21 | 18 | 38 | 36 | 39 |
| 3 | 5 | 42 | 37 | 32 |
| 9 | 5 | 42 | 41 | 37 |
| 18 | 15 | 42 | 43 | 40 |
| 15 | 20 | 30 | 40 | 41 |
| 10 | 18 | 36 | 28 | 32 |
| 8 | 15 | 31 | 34 | 30 |
| 14 | 10 | 37 | 38 | 39 |
| 15 | 13 | 36 | 35 | 41 |
| 19 | 22 | 36 | 34 | 39 |
| 15 | 10 | 41 | 32 | 43 |
| 12 | 6 | 45 | 39 | 36 |
| 26 | 20 | 31 | 41 | 43 |
| 25 | 12 | 38 | 40 | 40 |
| 21 | 27 | 40 | 27 | 47 |
| 23 | 14 | 41 | 40 | 42 |
| 19 | 18 | 35 | 42 | 48 |
| 18 | 20 | 37 | 35 | 33 |
| 19 | 17 | 36 | 34 | 30 |

| Total Trial 4 | % alive Trial 1 | % alive Trial 2 | % alive Trial 3 | % alive Trial 4 |
| --- | --- | --- | --- | --- |
| 37 | 40 | 47.22222222 | 58.53658537 | 59.45945946 |
| 41 | 42.10526316 | 40.74074074 | 38.46153846 | 51.2195122 |
| 36 | 60.71428571 | 56.25 | 54.28571429 | 38.88888889 |
| 33 | 40.90909091 | 46.15384615 | 45 | 48.48484848 |
| 42 | 54.54545455 | 32.43243243 | 55.17241379 | 26.19047619 |
| 27 | 45.94594595 | 41.02564103 | 50 | 44.44444444 |
| 41 | 60 | 66.66666667 | 58.82352941 | 63.41463415 |
| 33 | 44.44444444 | 52.38095238 | 48.57142857 | 51.51515152 |
| 33 | 38.46153846 | 33.33333333 | 51.06382979 | 45.45454545 |
| 37 | 75 | 75.86206897 | 48.57142857 | 51.35135135 |
| 32 | 45.94594595 | 61.29032258 | 62.16216216 | 53.125 |
| 32 | 54.28571429 | 51.72413793 | 41.66666667 | 46.875 |
| 39 | 48.71794872 | 51.72413793 | 50 | 51.28205128 |
| 39 | 63.15789474 | 50 | 51.28205128 | 53.84615385 |
| 8 | 64.28571429 | 62.16216216 | 59.375 | 37.5 |
| 14 | 61.9047619 | 85.36585366 | 70.27027027 | 64.28571429 |
| 33 | 42.85714286 | 60.46511628 | 40 | 54.54545455 |
| 35 | 60 | 67.5 | 48.7804878 | 42.85714286 |
| 28 | 38.88888889 | 50 | 43.75 | 35.71428571 |
| 23 | 64.51612903 | 52.94117647 | 56.66666667 | 34.7826087 |
| 24 | 40.54054054 | 50 | 41.02564103 | 58.33333333 |
| 28 | 47.22222222 | 60 | 51.2195122 | 53.57142857 |
| 41 | 61.11111111 | 58.82352941 | 56.41025641 | 46.34146341 |
| 25 | 56.09756098 | 43.75 | 51.1627907 | 60 |
| 18 | 46.66666667 | 53.84615385 | 41.66666667 | 66.66666667 |
| 46 | 74.19354839 | 58.53658537 | 48.8372093 | 56.52173913 |
| 37 | 65.78947368 | 72.5 | 55 | 67.56756757 |
| 48 | 55 | 62.96296296 | 46.80851064 | 43.75 |
| 37 | 48.7804878 | 55 | 35.71428571 | 62.16216216 |
| 37 | 62.85714286 | 40.47619048 | 62.5 | 51.35135135 |
| 38 | 45.94594595 | 34.28571429 | 51.51515152 | 47.36842105 |
| 36 | 44.44444444 | 50 | 46.66666667 | 52.77777778 |

|  | Normalized Survival |  |  |  | relative Difference |
| --- | --- | --- | --- | --- | --- |
| 51.3045668 | 0.77965769 | 0.92042922 | 1.14096247 | 1.15895062 | 0.22034231 |
|  | 0.820692305 | 0.7940958 | 0.74967086 | 0.99834216 | 0.17930769 |
|  | 1.183408994 | 1.09639363 | 1.05810686 | 0.75800053 | -0.183409 |
|  | 0.797377183 | 0.89960503 | 0.8771149 | 0.94503962 | 0.20262282 |
|  | 1.063169577 | 0.63215488 | 1.07538992 | 0.51049015 | -0.0631696 |
|  | 0.895552752 | 0.79964891 | 0.97457211 | 0.86628632 | 0.10444725 |
|  | 1.169486535 | 1.29942948 | 1.14655543 | 1.23604268 | -0.1694865 |
|  | 0.866286322 | 1.02098031 | 0.94672719 | 1.0041046 | 0.13371368 |
|  | 0.749670856 | 0.64971474 | 0.99530769 | 0.88597465 | 0.25032914 |
|  | 1.461858168 | 1.47866114 | 0.94672719 | 1.0009119 | -0.4618582 |
|  | 0.895552752 | 1.19463678 | 1.21163019 | 1.03548287 | 0.10444725 |
|  | 1.058106865 | 1.00817805 | 0.81214343 | 0.91366136 | -0.0581069 |
|  | 0.949583084 | 1.00817805 | 0.97457211 | 0.99956114 | 0.05041692 |
|  | 1.231038458 | 0.97457211 | 0.99956114 | 1.0495392 | -0.2310385 |
|  | 1.253021287 | 1.21163019 | 1.15730438 | 0.73092908 | -0.2530213 |
|  | 1.206613091 | 1.66390361 | 1.36966891 | 1.25302129 | -0.2066131 |
|  | 0.835347525 | 1.17855232 | 0.77965769 | 1.06316958 | 0.16465248 |
|  | 1.169486535 | 1.31567235 | 0.95080206 | 0.83534752 | -0.1694865 |
|  | 0.758000532 | 0.97457211 | 0.8527506 | 0.69612294 | 0.24199947 |
|  | 1.257512403 | 1.03189988 | 1.10451506 | 0.67796321 | -0.2575124 |
|  | 0.790193605 | 0.97457211 | 0.79964891 | 1.1370008 | 0.2098064 |
|  | 0.920429217 | 1.16948653 | 0.99834216 | 1.04418441 | 0.07957078 |
|  | 1.191143693 | 1.14655543 | 1.09951725 | 0.90326196 | -0.1911437 |
|  | 1.09342237 | 0.8527506 | 0.99723658 | 1.16948653 | -0.0934224 |
|  | 0.909600638 | 1.0495392 | 0.81214343 | 1.29942948 | 0.09039936 |
|  | 1.446139263 | 1.14096247 | 0.95190764 | 1.10169021 | -0.4461393 |
|  | 1.282331727 | 1.41312956 | 1.07202932 | 1.31698934 | -0.2823317 |
|  | 1.072029324 | 1.22723896 | 0.91236538 | 0.8527506 | -0.0720293 |
|  | 0.950802061 | 1.07202932 | 0.69612294 | 1.21163019 | 0.04919794 |
|  | 1.22517637 | 0.78893933 | 1.21821514 | 1.0009119 | -0.2251764 |
|  | 0.895552752 | 0.66827802 | 1.0041046 | 0.92327884 | 0.10444725 |
|  | 0.866286322 | 0.97457211 | 0.90960064 | 1.02871501 | 0.13371368 |

|  |  |  |
| --- | --- | --- |
| 0.07957078 | -0.1409625 | -0.1589506 |
| 0.2059042 | 0.25032914 | 0.00165784 |
| -0.0963936 | -0.0581069 | 0.24199947 |
| 0.10039497 | 0.1228851 | 0.05496038 |
| 0.36784512 | -0.0753899 | 0.48950985 |
| 0.20035109 | 0.02542789 | 0.13371368 |
| -0.2994295 | -0.1465554 | -0.2360427 |
| -0.0209803 | 0.05327281 | -0.0041046 |
| 0.35028526 | 0.00469231 | 0.11402535 |
| -0.4786611 | 0.05327281 | -0.0009119 |
| -0.1946368 | -0.2116302 | -0.0354829 |
| -0.008178 | 0.18785657 | 0.08633864 |
| -0.008178 | 0.02542789 | 0.00043886 |
| 0.02542789 | 0.00043886 | -0.0495392 |
| -0.2116302 | -0.1573044 | 0.26907092 |
| -0.6639036 | -0.3696689 | -0.2530213 |
| -0.1785523 | 0.22034231 | -0.0631696 |
| -0.3156724 | 0.04919794 | 0.16465248 |
| 0.02542789 | 0.1472494 | 0.30387706 |
| -0.0318999 | -0.1045151 | 0.32203679 |
| 0.02542789 | 0.20035109 | -0.1370008 |
| -0.1694865 | 0.00165784 | -0.0441844 |
| -0.1465554 | -0.0995173 | 0.09673804 |
| 0.1472494 | 0.00276342 | -0.1694865 |
| -0.0495392 | 0.18785657 | -0.2994295 |
| -0.1409625 | 0.04809236 | -0.1016902 |
| -0.4131296 | -0.0720293 | -0.3169893 |
| -0.227239 | 0.08763462 | 0.1472494 |
| -0.0720293 | 0.30387706 | -0.2116302 |
| 0.21106067 | -0.2182151 | -0.0009119 |
| 0.33172198 | -0.0041046 | 0.07672116 |
| 0.02542789 | 0.09039936 | -0.028715 |

**Supplementary Table 4:** Raw data used in the lifespan analysis.

